# Patterns of *Klebsiella pneumoniae* bacteremic dissemination from the lung

**DOI:** 10.1101/2024.04.03.587919

**Authors:** Caitlyn L. Holmes, Katherine G. Dailey, Karthik Hullahalli, Alexis E. Wilcox, Sophia Mason, Bridget S. Moricz, Lavinia V. Unverdorben, Matthew K. Waldor, Michael A. Bachman

## Abstract

Bacteremia, a leading cause of death, generally arises after bacteria establish infection in a particular tissue and transit to secondary sites. Studying dissemination from primary sites by solely measuring bacterial burdens does not capture the movement of individual clones. Here, by barcoding *Klebsiella pneumoniae*, a leading cause of bacteremia, we tracked pathogen dissemination following pneumonia. Variability in organ bacterial burdens was attributable to two distinct dissemination patterns distinguished by the extent of clonal expansion in the lungs. In “metastatic” dissemination, bacterial clones underwent heterogeneous clonal expansion within the lung and the dominant clones spread to secondary organs. In “direct” dissemination, bacterial clones exited the lungs without clonal expansion, leading to lower burdens in systemic sites. We uncover bacterial and host factors that govern these two modes of dissemination. Our data reveal unexpected heterogeneity in the dynamics of *Klebsiella* bacteremia and define a new paradigm for understanding within-host bacterial dissemination.

## Introduction

Sepsis is a systemic inflammatory syndrome with high mortality rates caused by a dysregulated immune response to infection (1, 2). Sepsis is highly associated with bloodstream infections, ~90% of which are caused by bacteria (3-5). Gram-negative species are leading causes of bacteremia and are highly linked to antimicrobial resistance (AMR) (6). *Enterobacterales* have been repeatedly classified as pathogens of urgent concern (7, 8) and are the most common causes of Gram-negative bacteremia. *Klebsiella pneumoniae* is a leading pathogen in both Gram-negative bacteremia and deaths by AMR (3, 6, 9).

Gram-negative bacteremia pathogenesis involves three phases (10). First, pathogens infect initial sites. Second, bacteria disseminate across host barriers and gain bloodstream access. Third, pathogens sustain bacteremia by avoiding immune clearance mechanisms in filtering organs like the spleen and liver (11). Pneumonia and bloodstream infections are the two most common conditions associated with death due to AMR organisms (9), and the lung is a common site of infection leading to secondary bacteremia. To prevent the progression of pneumonia to bacteremia, dissemination mechanisms by which bacteria exit the lung must be defined.

Experimentally defining host-pathogen interactions during dissemination is difficult since this process often occurs simultaneously with the phases of initial site infection and bloodstream survival. Murine pneumonia models can identify bacterial or host factors required for lung fitness and can assess bloodstream fitness by measuring bacterial abundance in tissues like the spleen and liver. However, these data must be carefully interpreted, as decreases in pathogen abundance at secondary sites can either be due to lower fitness at that site or differential dissemination from the lungs. Models that do not involve primary site infections, such as a tail vein injection, define bacteremia fitness defects in a dissemination-independent manner. For example, we identified GmhB, involved in ADP-heptose biosynthesis for the lipopolysaccharide (LPS) inner core and a ligand for the pro-inflammatory sensor alpha-kinase 1 (ALPK1) (12, 13), as a conserved Gram-negative bacteremia fitness factor (14) in both pneumonia and tail-vein models. However, the defect of the *K. pneumoniae gmhB* mutant in the spleen was larger in the pneumonia model despite no defect in the lung, indirectly suggesting that gmhB may affect both dissemination and sustaining bacteremia. Thus, bacterial tissue burden can measure defects at the first and last bacteremia phases (initial site infection and avoiding immune clearance), but metrics to specifically quantify dissemination are lacking (10).

Isogenic barcoding of bacteria has been used to study infection dynamics in various models (15-27). This approach enables the quantitative tracking of individual bacterial clones during different phases of infection, including the measurement of infection bottlenecks and clonal replication within tissues. One approach for analyzing barcoded bacteria is STAMPR (Sequence-Tagged Analysis of Microbial Populations in R), which employs population genetics and resampling approaches to quantify bacterial expansion, infection bottlenecks, and dissemination patterns (23, 28). By comparing the presence and abundance of barcodes across sites relative to the inoculum, STAMPR can trace clones between tissues, enabling inferences regarding *in vivo* dissemination routes.

Here, we applied bacterial barcoding and STAMPR to a murine model of bacteremia originating from pneumonia. Initially, wild-type *K. pneumoniae* bacteremia dynamics were measured. Two distinct dissemination patterns termed “metastatic” and “direct” were identified and correlated with clonal expansion in the lung or absence thereof, respectively. Metastatic dissemination led to higher bacterial burden in systemic organs. STAMPR analyses using bacterial and host mutants were used to define how dynamics change after disruption of bacterial ADP-heptose biosynthesis (GmhB) or protein translocation (TatC), and host Nox2 NADPH oxidase (CybB) or monocyte chemokine receptor (Ccr2). Disruption of *gmhB* eliminated metastatic dissemination, leading to only direct dissemination and low spleen burdens. Disruption of Nox2 also yielded only direct dissemination, but with high spleen burdens likely due to attenuated host defenses. This study uncovers informative yet previously hidden patterns that define bacterial spread from the lung to the bloodstream and correlate with bacterial burdens at systemic sites. Furthermore, the framework for analysis of pathogen dissemination presented here should be broadly useful for studying bacteremia-associated infection dynamics.

## Results

### Bacterial barcoding reveals the dynamics of *K. pneumoniae* dissemination from pneumonia

To characterize *K. pneumoniae* infection dynamics, we barcoded the hypervirulent strain KPPR1 (29). This library, KPPR1-STAMPR, contained ~40,000 unique barcodes inserted at the Tn7 site. The 25-nucleotide barcodes did not influence bacterial fitness, and KPPR1-STAMPR had an even abundance of barcodes across the library (Supplemental Figure 1A-B). Deep sequencing the barcodes can be used to estimate the founding population, the number of individual clones from the inoculum to give rise to infection (N_s_, Online Methods). Smaller founding populations are indicative of tighter bottlenecks. To confirm that the library enabled accurate calculations of founding population, we created a standard curve using *in vitro* bottlenecks with serial dilutions, and verified the library is appropriate for estimating founding populations up to ~10^5^ (Supplemental Figure 1C).

To measure baseline dynamics of *K. pneumoniae* infection in a model of pneumonia that progresses to bacteremia (30), WT C57BL/6 mice were retropharyngeally inoculated with 1x10^6^ CFU KPPR1-STAMPR. Compared to the inoculum size, the *K. pneumoniae* population expanded in the lung ~10-100x (Figure 1A). Bacterial CFU recovery from the spleen and liver were typically lower than the inoculum size, indicating replication restriction within tissues or tight bottlenecks between primary and secondary sites. Bacterial abundance in the liver and spleen were variable but correlated with lung burden (Supplemental Figure 2). Based on the number of unique barcodes recovered from each tissue relative to the reference, the STAMPR pipeline estimated the founding population size through a resampling approach (N_s_, Online Methods). The lung contained a large founding population (Figure 1B) that produced the observed CFU, with an average N_s_ of ~3,850 (Supplemental Data 1). Secondary organs had much lower N_s_ values, with an average founding population of ~16, 40, and 60 clones in the spleen, liver, and blood, respectively, suggesting that the major bottleneck occurs between the lungs and systemic sites. Furthermore, since only ~0.7% of clones from the lung were identified at a secondary site, a tight bottleneck limits *K. pneumoniae* exit from the lung (Figure 1B).

**Figure 1.**
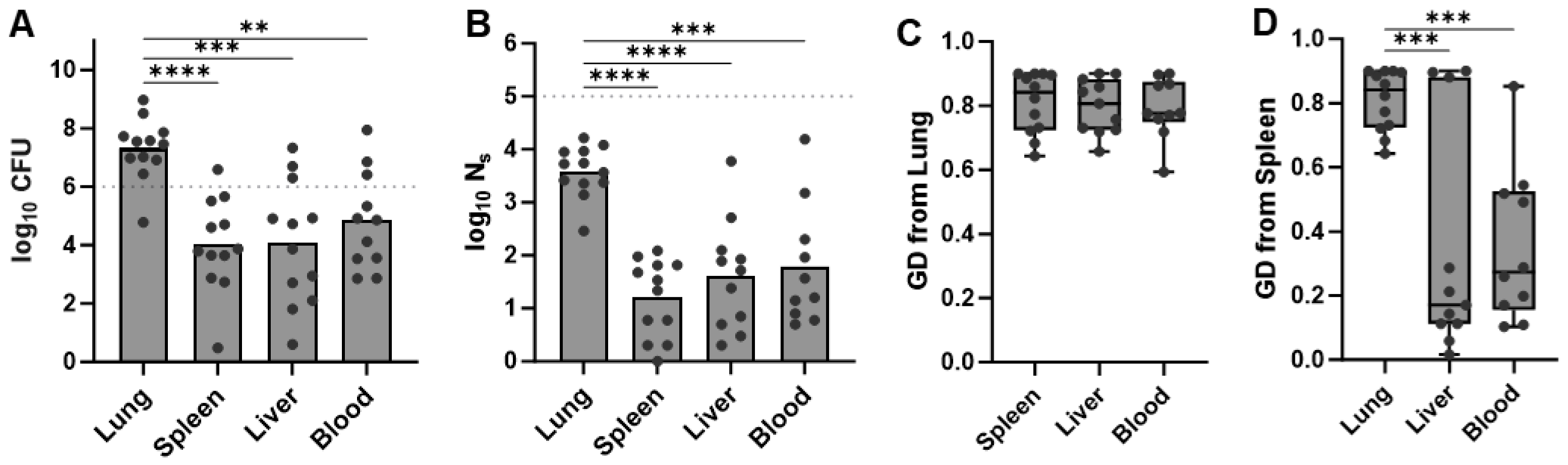
*K. pneumoniae* population dynamics during bacteremic pneumonia. Mice were infected with KPPR1-STAMPR, a wild-type *K. pneumoniae* barcoded library in a model of pneumonia that progresses to bacteremia. Tissues were harvested 24 hours post-infection and analyzed with quantitative culture and the STAMPR pipeline. (A) The bacterial burden in each tissue at the time of harvest is displayed as log_10_ CFU. The dotted line indicates the infection inoculum, 1x10^6^ CFU/mouse. (B) The founding population size in each tissue was estimated by the STAMPR pipeline and is displayed as log_10_ N_s_. The dotted line indicates the resolution limit of the input library, ~1x10^5^ N_s_, which is the maximum potential complexity for any tissue. The genetic distance between the lung and secondary organs (C) or the spleen and other secondary organs (D) was modeled by the STAMPR pipeline. For all, \*\**p*<0.01, \*\*\**p*<0.001, \*\*\*\**p*<0.0001 by ordinary one-way ANOVA with Tukey’s multiple comparisons test; n=12 mice in 4 independent trials. STAMPR analysis was excluded for one mouse with no detectable blood CFU and any sample with low sequencing quality.

We next evaluated the extent of barcode similarity between sites using genetic distance (GD). High GD values indicate dissimilarity between groups. Low GD values indicate similarity, due to either sharing many barcodes or sharing a few highly abundant barcodes. The GD between the lung and secondary sites was high, indicating the lung was substantially dissimilar from the spleen, liver, and blood (Figure 1C). GDs between the spleen and other secondary sites were lower, demonstrating that systemic sites were more similar to each other than the lung (Figure 1D). Thus, after bacteria enter circulation, there is a greater likelihood of spread between secondary organs or an initial seeding event of the same clones to these sites. However, GD between secondary organs was not always uniform as some liver samples were very dissimilar from corresponding spleens (Figure 1D).

### Two distinct modes of *K. pneumoniae* dissemination from the lungs

The frequency of each barcode in individual tissues can be displayed as a frequency plot (Figure 2A-D). Analysis of lung and spleen frequency plots revealed two distinct patterns of barcode sharing between these sites. In one subset of mice (5/12), there were clearly dominant (highly abundant) clones in the lung (Figure 2A, red circles), and these were also the dominant barcodes in the spleen (Figure 2B). In a second subset (7/12), there was little variation in clonal abundance (more evenly distributed barcodes) across the population of bacteria in the lung; the most abundant clones in the lung (Figure 2C, red circles) were not typically abundant in the spleen (Figure 2D, red circles).

**Figure 2.**
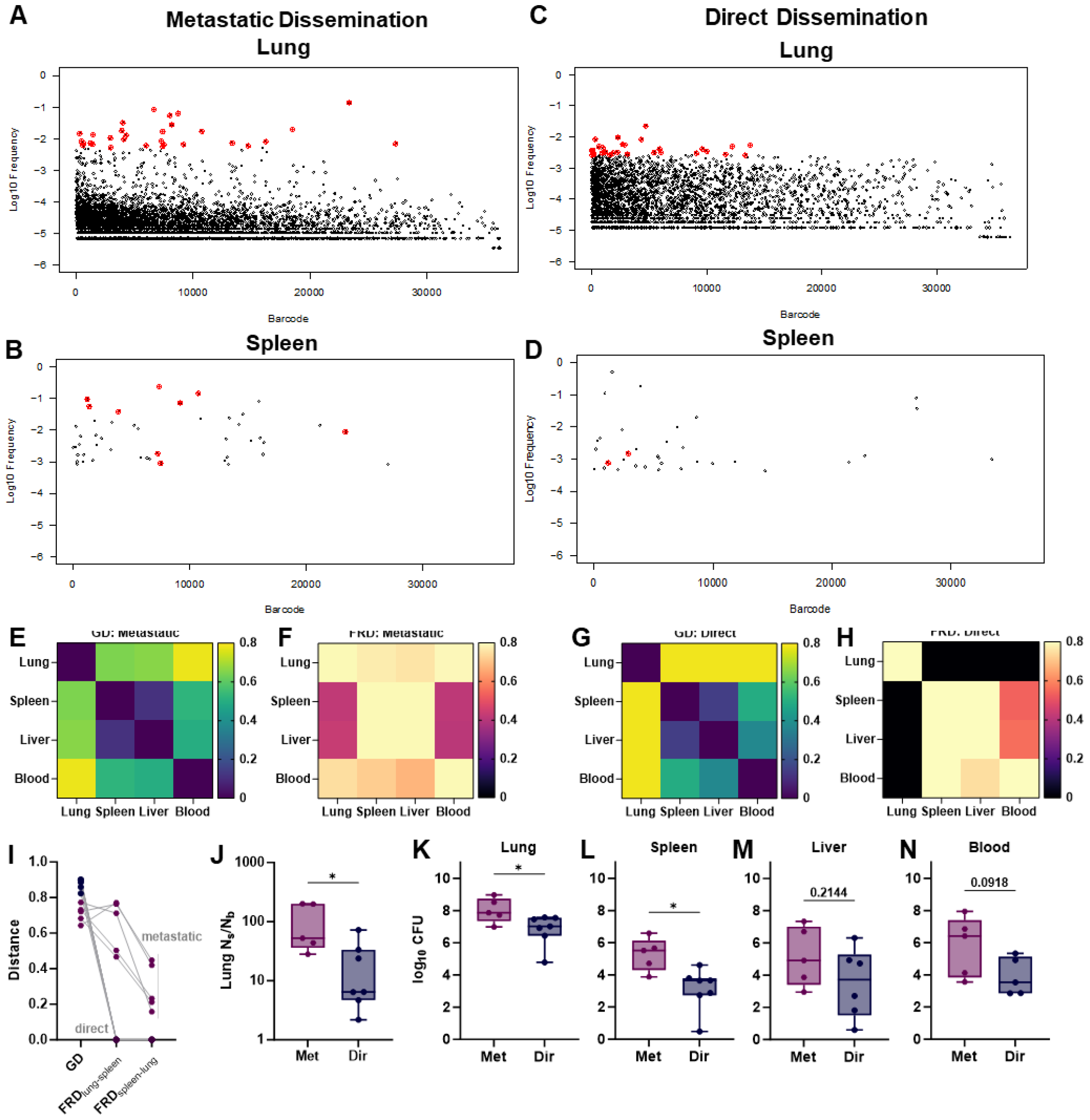
*K. pneumoniae* dissemination from the lung occurs in two distinct patterns. Mice were infected with a wild-type *K. pneumoniae* barcoded library in a model of bacteremia originating from pneumonia. The STAMPR pipeline calculated the frequency of individual barcodes within the lung and spleen of each mouse. (A-D) Unique 25-nt barcodes were assigned a random tag and plotted on the x-axis, the log_10_ frequency of each barcode within the indicated tissue is plotted on the y-axis. The most abundant barcodes within the lung for each mouse are highlighted in red, and indicated in the spleen if the barcode was also found at that site. During metastatic dissemination, the top thirty most abundant barcodes in the lung (A) replicated more than other clones, and the same clones were more likely to be found in the corresponding spleen (B). During direct dissemination, the most abundant barcodes in the lung did not have extensive replication beyond other barcodes (C) and were less likely to be found in the corresponding spleen (D). Using the STAMPR pipeline, genetic distance (GD; E,G) and fractional genetic distance (FRD; F, H) were calculated to determine the barcode similarity and sharing between sites. Representative GD and FRD heat maps between the lung, spleen, liver, and blood for two individual mice that demonstrated metastatic (E,F) or direct (G,H) patterns of dissemination are displayed. Individual GD and FRD between the lung and spleen or the spleen and lung are displayed for each mouse (I), with indications for defining dissemination patterns as metastatic of direct. The N_s_/N_b_ ratio (J) reflects the evenness of lung expansion for animals experiencing metastatic of direct. Bacterial burden, displayed at log_10_ CFU, is compared across the (K) lung, (L) spleen, (M) liver, or (N) blood for each mouse demonstrating metastatic or direct dissemination. For (I-N), n=12 mice infected across four independent trials; \**p*<0.05 by unpaired *t-*test. STAMPR analysis was excluded for one mouse with no detectable blood CFU, and any sample with low sequencing quality.

A deeper analysis of the relatedness of barcodes between the lung and spleen further distinguished the two patterns of *K. pneumoniae* lung dissemination. Overall, the GD was high between the lung and each site of dissemination (spleen, liver, blood; Figure 1C), but frequency plots indicated some sharing of barcodes with the lung. To further examine this sharing, a fractional genetic distance metric (FRD_x-y_) was used to quantify the fraction of clones shared between site x and y relative to the total number of clones in y. For example, a high FRD_lung-spleen_ indicates that their shared barcodes are a high proportion of the total spleen barcodes. A corresponding lower FRD_spleen-lung_ would suggest the lung is a large reservoir of clones, only some of which are shared with the spleen. By measuring FRD_lung-spleen_, we observed two distinct patterns of clonal sharing (Figure 2E-H). In one subset of mice, there were many clones shared between the lung and spleen (high FRD_lung-spleen_, Figure 2E, F), even though the lungs and spleens were dissimilar overall (high GD, Figure 1C). Dissimilar lungs and spleens with many shared clones can be explained by uneven expansion of clones in the lung that are present in the spleen at distinct frequencies (Figure 2A-B). To quantify uneven replication, we plotted N_s_/N_b_ ratios (Figure 2I). N_b_ is a STAMPR metric that calculates founding population sizes based on the frequency of barcodes, rather than the number of unique barcodes (N_s_). When barcode abundances are relatively even, N_s_ and N_b_ ratios are close to one. During uneven bacterial replication, N_b_ will decrease but N_s_ will remain constant, and thus the N_s_/N_b_ ratio will increase (23). Mice with high FRD_lung-spleen_also had high N_s_/N_b_ ratios. Furthermore, they had significantly elevated bacterial burdens in the lungs and spleens, with trends toward increased burden in the liver and blood (Figure 2J-N). We define this pattern as “metastatic dissemination,” in which clonal expansion in one organ is associated with translocation to another.

In a second subset, lung clones were not abundant in the spleen (FRD_lung-spleen_=0; Figure 2G-H). Individual barcode frequency and N_s_/N_b_ ratios confirmed that these animals had more even expansion in the lung (Figure 2C,D,J). In this group, clones from the inoculum translocated to the spleen without substantial expansion in the lungs, which we termed “direct dissemination”. This pattern was associated with lower bacterial burdens in the lung and spleen (Figure 2K, L). Together, these data reveal two dissemination patterns that were not apparent from CFU alone.

### Metastatic dissemination is dependent on *K. pneumoniae* GmhB and host Nox2

We next sought to determine how *K. pneumoniae* fitness factors that differ in their importance for primary site infection and bloodstream survival affect patterns of *K. pneumoniae* dissemination from the lung. Barcoded libraries were generated in Δ*gmhB* and Δ*tatC* mutants, named *gmhB-*STAMPR and *tatC-*STAMPR, respectively. TatC is part of the twin-arginine transporter required for moving multiple folded protein substrates across the cytoplasmic membrane (31). Unlike GmhB, which is dispensable in the lung, TatC is required for both primary and secondary site fitness during bacteremia in multiple species (32, 33). Based on bacterial burden, it is unclear if either is involved in dissemination.

Wild-type mice were infected with *gmhB-*STAMPR and *tatC*-STAMPR using the pneumonia model, and dissemination metrics were quantified. The mutant libraries had an even abundance of barcodes (Supplemental Figure 1D-E). To determine if GmhB and TatC influence lung dissemination, GD and FRD between the lung and spleen or lung and liver were calculated. GmhB was required for metastatic dissemination as all mice infected with *gmhB*-STAMPR exhibited direct dissemination from the lung to the spleen and liver (Figure 3A-B). TatC also influenced dissemination, but to a lesser degree, as 1/5 mice and 3/5 mice displayed metastatic dissemination from the lung to the spleen and liver, respectively (Figure 3C-D), compared to 5/12 displaying metastatic dissemination (lung to spleen) following infection with KPPR1-STAMPR (Figure 2I-N). Each mouse infected with KPPR1-STAMPR and *gmhB-*STAMPR demonstrated the same dissemination pattern between the lung and spleen as the lung and liver (Supplemental Figure 3A-C). In contrast, two *tatC-*STAMPR-infected mice displayed direct dissemination to the spleen but metastatic dissemination to the liver (Supplemental Figure 3D-F). Thus, GmhB is required for metastatic dissemination from the lung, whereas *tatC* may influence dissemination to specific sites.

**Figure 3.**
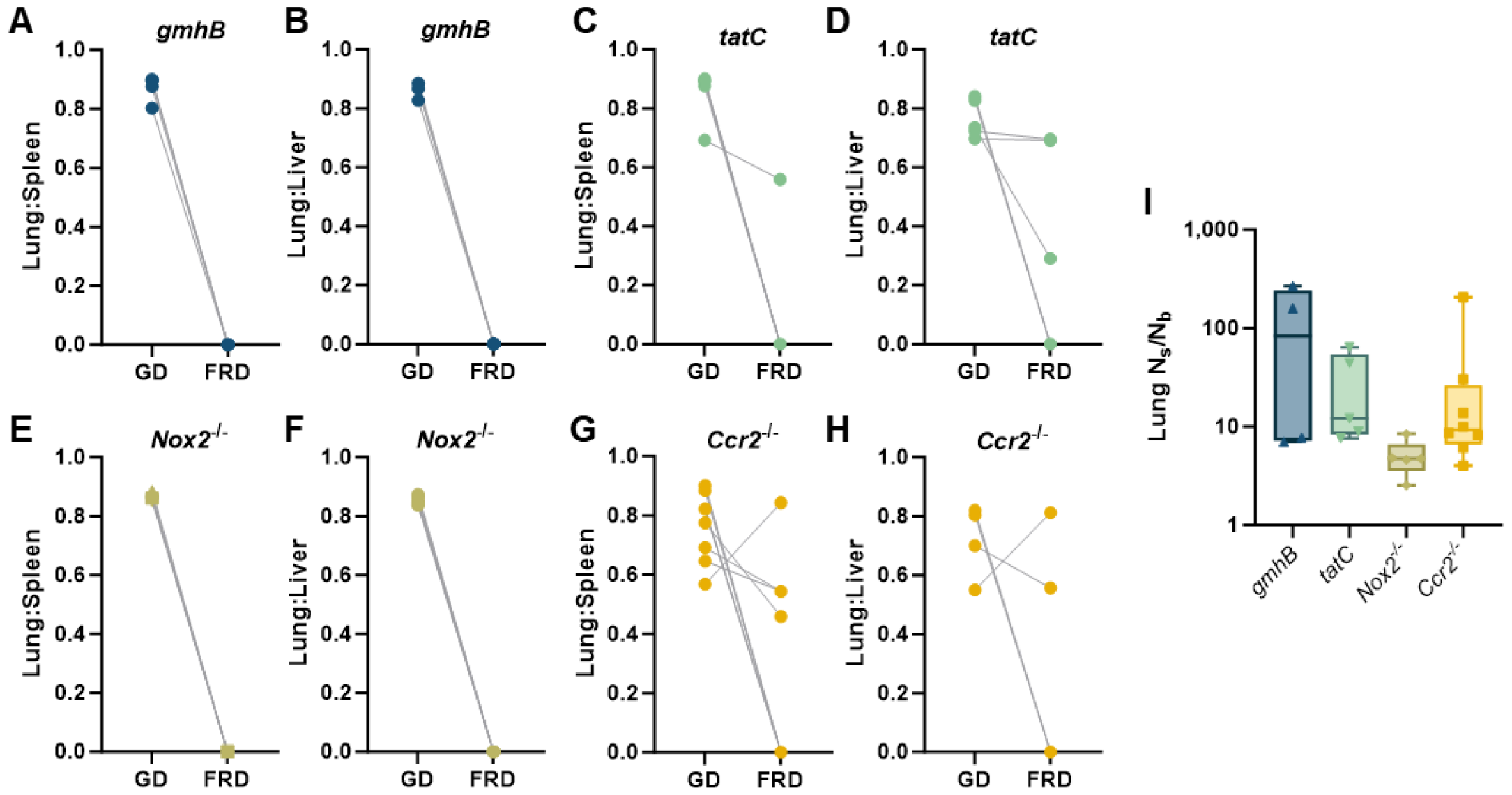
Metastatic dissemination is dependent on bacterial GmhB and host Nox2. Wild-type mice were infected with a (A,B) *gmhB*-STAMPR or (C,D) *tatC*-STAMPR *K. pneumoniae* barcoded libraries, while (E,F) *Nox2*^*-/-*^ or (G,H) *Ccr2*^*-/-*^ mice were infected with the wild-type KPPR1-STAMPR library. Genetic distance (GD) or fractional genetic distance (FRD) was calculated to determine genetic similarity and the fraction of shared clones between the lung and spleen (A, C, E, G) and the lung and liver (B, D, F, H). The lung N_s_/N_b_ was calculated for all group (I). All animals were inoculated retropharyngeally with 1x10^6^ CFU/mouse and tissues were harvested 24 hours post infection. N=4-8 mice in two independent trials; in (A-H), lines connect GD and FRD from the same mouse. STAMPR analysis was excluded for any sample with low sequencing quality.

To determine how host defenses determine the patterns of dissemination, we performed STAMPR analysis of wild-type *K. pneumoniae* in mice defective in NADPH oxidase Nox2 (*Cybb*^*-/-*^), and the monocyte chemokine receptor Ccr2 (*Ccr2*^*-/-*^), which are required for the phagocyte oxidative burst and monocyte recruitment, respectively. Nox2 restricts *K. pneumoniae* replication in the lung but has more subtle influences in the spleen and liver (32). After lung infection, all *Nox2*^*-/-*^ mice exhibited direct dissemination, suggesting that phagocyte NADPH oxidase is required for the metastatic pattern of *K. pneumoniae* dissemination (Figure 3E-F). Since Nox2 is involved in oxidative bursts across neutrophils and monocytes, which are efficiently recruited during *K. pneumoniae* lung infection (14), we sought to resolve which immune cell subset may influence *K. pneumoniae* dissemination by infecting *Ccr2*^*-/-*^ mice with KPPR1-STAMPR. Both metastatic and direct dissemination were observed in *Ccr2*^*-/-*^ mice, indicating that monocytes are dispensable for metastatic dissemination (Figure 3G-H). As with the *tatC-*STAMPR library, KPPR1-STAMPR infections in the *Ccr2*^*-/-*^ mice had mixed dissemination patterns. In 2/6 mice, metastatic dissemination to the spleen and direct dissemination to the liver were observed. Similar to TatC, Ccr2 may influence dissemination patterns to specific sites but is not required for metastatic dissemination. To further assess the association between the dynamics of lung expansion and the mode of dissemination, N_s_/N_b_ was calculated (Figure 3I). *Nox2*^*-/-*^ had even expansion, with consistently low N_s_/N_b_, whereas the *gmhB* and *tatC* mutants, and *Ccr2*^*-/-*^ mice had more variation. Together, these data demonstrate that host Nox2 influences bacterial lung dissemination in a monocyte-independent manner. Increased metastatic dissemination in the absence of Nox2 suggest that oxidative bursts may impose stresses that result in heterogenous *K. pneumoniae* replication in the lungs.

### Patterns of dissemination from the lung influence the similarity of populations in the liver and spleen

To obtain a broader perspective of *K. pneumoniae* infection dynamics, the similarities of barcodes in different organs were analyzed across all experiments (Figure 4A-C). Barcodes found in the lungs were largely distinct from secondary sites regardless of the *K. pneumoniae* strain or mouse genotype (Figure 4A-B). KPPR1-infected wild-type mice with metastatic dissemination had significantly higher similarity (lower GD) between the lung and secondary organs than mice with direct dissemination, including the *gmhB* or *Nox2*^*-/-*^ groups (Figure 4A-B). While the lung was largely distinct from secondary organs, clonal sharing between secondary sites varied with the mode of dissemination. Splenic and hepatic populations were similar in KPPR1-infected wild-type mice with metastatic dissemination. Clonal sharing between the spleen and liver in animals with direct dissemination varied, with some animals having similar clones and others having distinct clones (Figure 4C). These observations reveal that the extent of replication in the lungs, which can result in metastatic or direct dissemination modes, can influence the genetic similarity of bacteria between systemic sites. Defining dissemination modes from the primary site is therefore critical for interpreting bacterial dissemination between distinct secondary sites.

**Figure 4.**
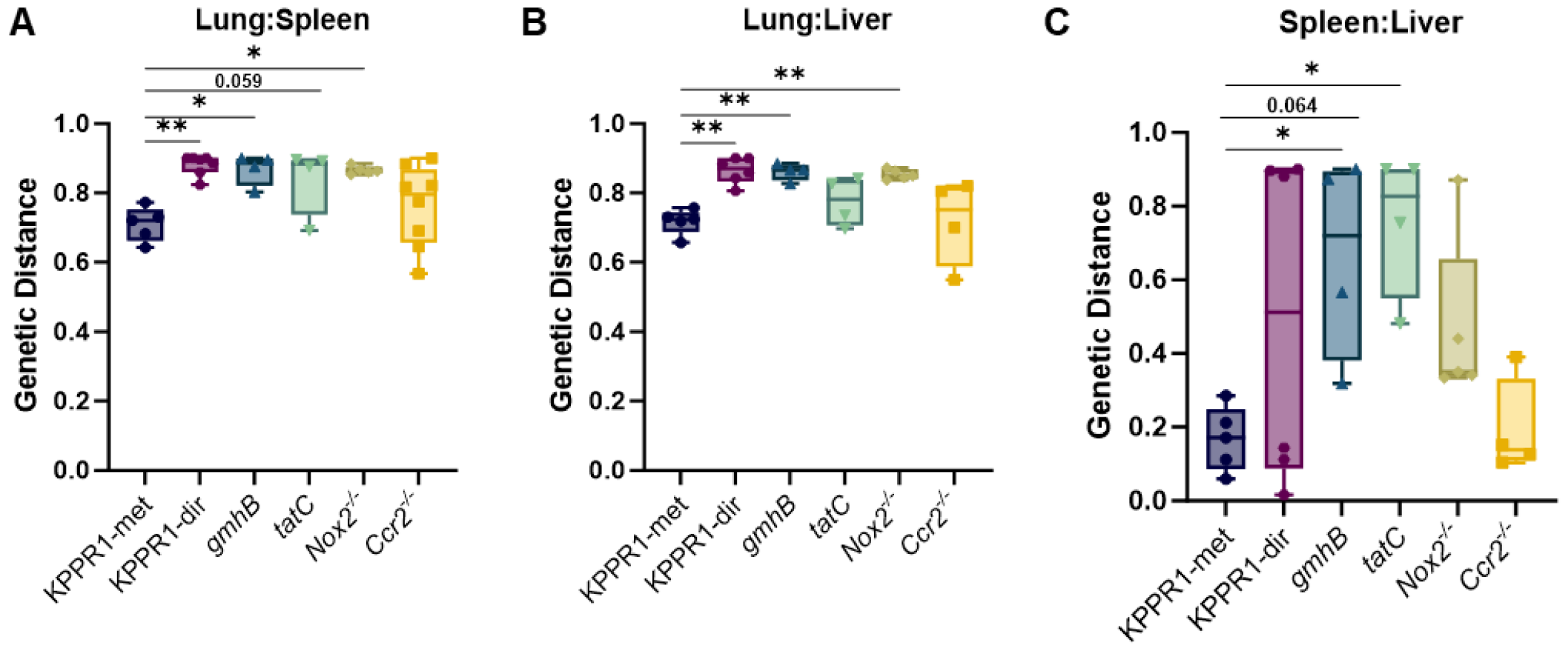
Secondary organs are genetically distinct compartments during bacteremia. Wild-type mice were infected with a KPPR1-STAMPR, *gmhB*-STAMPR, or *tatC*-STAMPR *K. pneumoniae* barcoded library. *Nox2*^*-/-*^ or *Ccr2*^*-/-*^ mice were infected with the KPPR1-STAMPR barcoded library. In all experiments, an inoculum of 1x10^6^ CFU/mouse was administered in a model of pneumonia and tissues were harvested 24 hours post-infection. Wild-type mice infected with the KPPR1 library were divided into groups demonstrating metastatic (KPPR1-met) or direct (KPPR1-dir) dissemination as defined in Figure 2. The genetic distance between the (A) lung to spleen, (B) lung to liver, and (C) spleen to liver were evaluated by the STAMPR pipeline. For all, n=4-8 mice in at least two independent trials; \**p*<0.05 and \*\**p*<0.01 using an ordinary ANOVA with Dunnet’s multiple comparisons test comparing each group to the KPPR1-met group.

### *K. pneumoniae* population dynamics in secondary sites are influenced by the mode of lung dissemination

To understand how host and bacterial factors control infection bottlenecks and bacterial replication, we quantified bacterial burden (Figure 5A-D), founding populations (Figure 5E-H), and bacterial replication across all sites (Figure 5I-L). Although metastatic dissemination (KPPR1-met) was generally associated with higher bacterial burdens (Figure 2K-N), founding populations in secondary sites were not significantly different (Figure 5E-H). Instead, mice with direct dissemination (KPPR1-dir) trended towards lower expansion per founder across sites (CFU/N_s_, Figure 5I-L). Thus, metastatic dissemination is associated with more apparent clonal expansion in secondary sites, even though host bottlenecks between the lung and secondary sites are similar.

**Figure 5.**
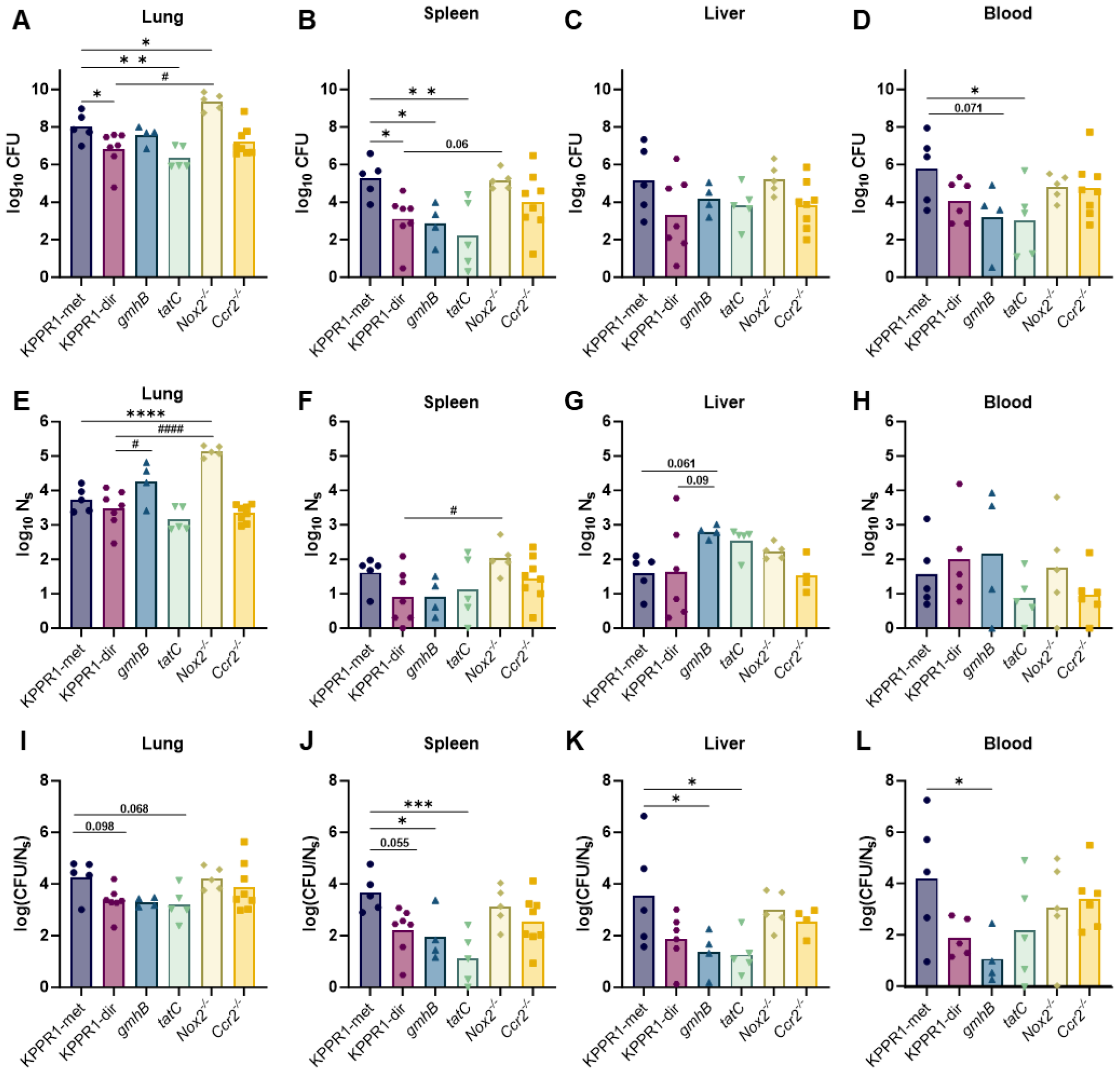
*K. pneumoniae* dynamics in secondary organs is influenced by the mode of dissemination. Wild-type mice were infected with a KPPR1-STAMPR, *gmhB*-STAMPR, or *tatC*-STAMPR *K. pneumoniae* barcoded library. *Nox2*^*-/-*^ or *Ccr2*^*-/-*^ mice were infected with the KPPR1-STAMPR barcoded library. For all, an inoculum of 1x10^6^ CFU/mouse was administered in a model of pneumonia and tissues were harvested 24 hours post infection. Wild-type mice infected with the KPPR1 library were divided into groups demonstrating metastatic (KPPR1-met) or direct (KPPR1-dir) dissemination as defined in Figure 2. The bacterial burden at the time of harvest in the (A) lung, (B) spleen, (C) liver, or (D) blood for each infection group is displayed as log_10_ CFU. The founding population in each tissue was estimated by the STAMPR pipeline for the (E) lung, (F) spleen, (G) liver, or (H) blood for each group and displayed as log_10_ N_s_. A measure of total CFU per founder is represented by log_10_(CFU/N_s_) for the (I) lung, (J) spleen, (K) liver, or (L) blood for each group. For all groups, n=4-12 mice in at least two independent trials. \**p*<0.05, \*\**p*<0.01, \*\*\**p*<0.001, \*\*\*\**p*<0.0001 using an ordinary ANOVA with Dunnet’s multiple comparisons test comparing each group to the KPPR1-met group; ^#^*p*<0.05, ^####^*p*<0.0001 using an ordinary ANOVA with Dunnet’s multiple comparisons test comparing each group to the KPPR1-dir group.

As previously described, GmhB was dispensable for fitness in the lung but had a significant spleen fitness defect (Figure 5A-D) and was required for metastatic dissemination (14). Despite both having direct dissemination, there was a larger *gmhB* founding population in the lung than observed with KPPR1-dir (Figure 5E). Although the number of *gmhB* founders in the spleen was similar to that found with wild-type bacterial infections (Figure 5F), the CFU was 100-fold lower than KPPR1-met (Figure 5B), indicating that expansion (CFU/N_s_) was lower in the *gmhB* mutant (Figure 5J). These observations are consistent with published findings that *gmhB* is defective in the spleen *ex vivo* and *in vivo* (14). In the blood, *gmhB* CFU was lower than that of KPPR1-met (Figure 5D); lower CFU is attributed to reduced expansion (Figure 5L) rather than tighter infection bottlenecks (Figure 5H, similar N_s_). In contrast, *gmhB* experienced wider bottlenecks in the liver than KPPR1-met (Figure 5G, higher N_s_) but had equivalent CFU (Figure 5C), indicating significantly less expansion than KPPR1-met (Figure 5K, lower CFU/N_s_). Together, these observations demonstrate that GmhB is required for metastatic dissemination and enhances splenic fitness but has a distinct function in the liver. Direct dissemination of *gmhB* did not result in fewer founders at secondary sites compared to WT direct dissemination, suggesting that GmhB primarily influences expansion in the lung. These findings reveal complex, tissue-specific functions of GmhB, and in its absence, dissemination patterns and replication are altered across secondary sites.

In contrast to GmhB, TatC was required for fitness in the lung, spleen, and blood as evidenced by lower CFU at all sites (Figure 5A-D). However, reductions in tissue burdens were not due to a difference in founding populations (Figure 5E-H). As with GmhB, there was a trend toward elevated numbers of *tatC* founders in the liver but this did not reach statistical significance (*p*=0.144). Rather, TatC was required for expansion in the spleen and liver (Figure 5J-K) and trended towards less expansion in the lung (Figure 5I). Thus, TatC is dispensable for exiting the lung and reaching systemic sites but is instead required for expansion (lower CFU/N_s_ values). Alternatively, reductions in CFU/N_s_ values could reflect a reduction in reseeding of the same clones from the lung to the spleen. In the lung, lower CFU/N_s_ likely represents a defect in replication.

Mice lacking Nox2 displayed significantly elevated lung CFU compared to wild-type mice (Figure 5A). This correlated to many more *K. pneumoniae* founders in the lungs of *Nox2*^-/-^ mice than KPPR1-met or -dir, indicating defective host clearance and a wider bottleneck in the lung (Figure 5E). Although *Nox2*^-/-^ mice experienced direct dissemination, bacterial burdens in the liver, spleen, and blood of *Nox2*^*-/-*^ mice were similar to KPPR1-met mice (Figure 5B-D), with similar founding population sizes (Figure 5F-H) and expansion of founders (Figure 5J-L) across sites. *Nox2*^*-/-*^ mice displayed even replication in the lung yet had a larger number of founders (Figure 3I). Thus, although Nox2 tightens infection bottlenecks in the lung, oxidative bursts may unveil underlying heterogeneity in *K. pneumoniae* replication in the lung. In contrast, mice lacking Ccr2 had similar bacterial burden, founding populations, and expansion across sites (Figure 5A-L) as wild-type mice with metastatic dissemination. Thus, monocytes are likely dispensable for controlling *K. pneumoniae* expansion and determining modes of dissemination during early bacteremia. Together, these analyses reveal how perturbations in host and bacterial factors influence tissue-specific *K. pneumoniae* replication and dissemination.

## Discussion

Here, using a murine model of bacteremic pneumonia, we leveraged barcoded *K. pneumoniae* to investigate bacterial dissemination from the lung. This second phase of bacteremia pathogenesis has been impossible to study independently from other phases using tissue CFU alone. Using high-complexity libraries of barcoded bacteria with varied bacterial and mouse genotypes, we defined two modes of lung dissemination and identified bacterial and host factors that influence each. Metastatic dissemination was associated with higher tissue CFU and uneven clonal expansion, and dominant lung clones were highly abundant in the spleen. Direct dissemination was associated with lower tissue CFU, and the lung and spleen had distinct populations (Figure 6). The bacterial factor GmhB was required for metastatic dissemination, and the direct dissemination of *gmhB* was associated with less severe infection. The host defense effector Nox2 was required for both metastatic dissemination and control of the initial infecting population, leading to severe infection with high systemic burdens in *Nox2*^-/-^ mice. These findings uncover a critical role for replication patterns at the primary site of infection in modulating dissemination. We propose that uncovering the hidden variables that influence dissemination will reveal principles of infection underlie disease severity during pneumonia and progression to bacteremia.

**Figure 6.**
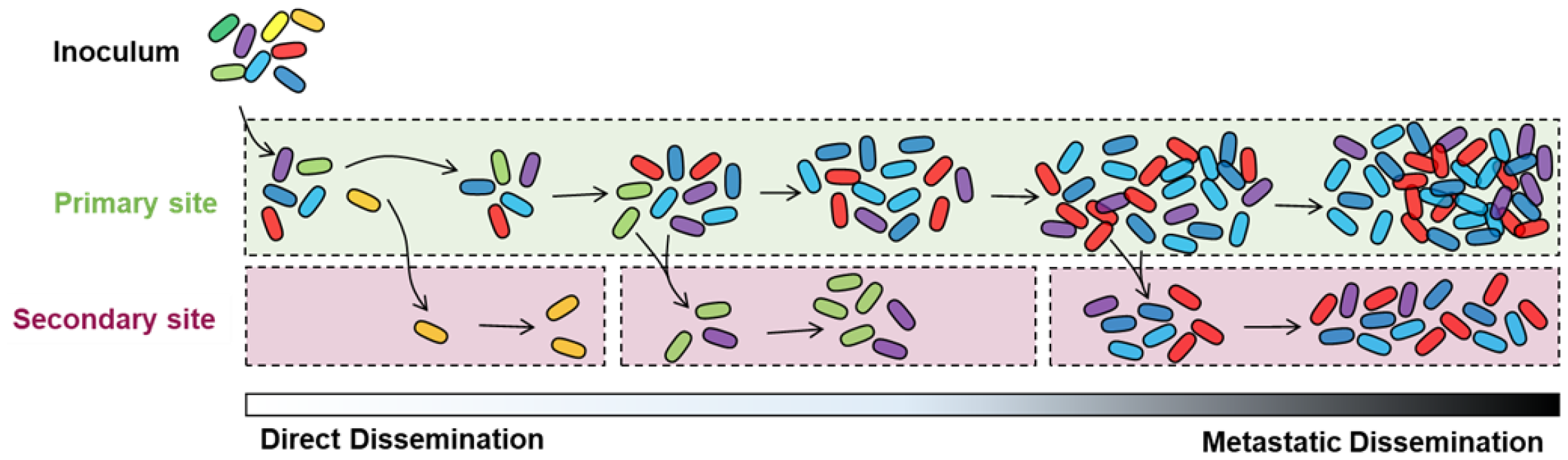
Bacterial barcoding reveals two patterns of lung dissemination during early *K. pneumoniae* bacteremia. Bacterial barcoding and STAMPR analysis revealed infection dynamics that are unresolved with quantitative culture and CFU values alone. In direct dissemination, replication is not necessary prior to translocation from the initial site. This pattern may arise due to defects in the host barrier integrity during infection. In metastatic dissemination, clonal replication in the lung leads to the presence of dominant clones in secondary sites. It is likely that bacterial factors influencing replication ability at the initial site also influence the ability for metastatic dissemination to occur. Metastatic dissemination may be influenced by time, where clones which have more time to replicate are more likely to translocate and be identified at a secondary site. While two patterns of dissemination are described here, these modes likely represent a spectrum of translocation dynamics.

The primary distinguishing feature between metastatic and direct dissemination is the extent to which bacterial replication promotes the transit of clones from primary to secondary compartments (Figure 6). Metastatic dissemination, here detectable with high FRD values, involves clones prominent in the lung population disseminating to the blood, liver, and spleen. This metastatic pattern suggests that clones that survive initial host defenses replicate in the lung and eventually disseminate. Fitness factors that enable lung survival and replication are predicted to be required for metastatic dissemination. Importantly, the similarity between the lung and other sites could be explained by a single dissemination event with expansion in the second site, or repeated dissemination of dominant lung clones. The correlation of high FRDs with uneven clonal replication in the lung and elevated CFU in the spleen suggests that other processes, which can be assessed with STAMPR-based metrics, such as N_s_/N_b_, underlie the extent of metastatic dissemination. In contrast, direct dissemination occurs when bacteria do not undergo clonal expansion in the initial site. Several biological mechanisms may explain how direct dissemination occurs. One is that a small degree of transit occurs early following infection, or even during the inoculation process, where organs are seeded with different subpopulations of clones that progress on different trajectories of survival and replication. Alternatively, it is possible that low-abundance clones occupy privileged niches in the primary site that enable these clones to disseminate more efficiently.

Barcoding provided insights on the bacteremia fitness factors GmhB and TatC that could not be elucidated from CFU counts alone (14). GmhB is dispensable for lung fitness but required for fitness in the spleen and liver. Our data demonstrates that GmhB also influences modes of *K. pneumoniae* dissemination. GmhB is involved in the biosynthesis cascade of ADP-heptose, which can be integrated into the LPS core or secreted and sensed as a soluble inflammatory mediator by cytosolic ALPK1 (13, 14, 34). It remains unknown which ADP-heptose function is the most relevant for influencing modes of *K. pneumoniae* dissemination from the lung. Our study also revealed new information about TatC, which may influence dissemination patterns to individual secondary organs in addition to enhancing primary site fitness.

Nox2 is required for metastatic dissemination independent of inflammatory monocytes. It is therefore likely that metastatic dissemination is dependent on other immune subsets, like neutrophils or alveolar macrophages. Since metastatic dissemination is marked by high bacterial replication and lung CFU, it is curious that this pattern requires oxidative bursts from immune cells, which is typically associated with bacterial clearance. Neutrophil killing of *K. pneumoniae* can be inhibited by soluble mediators secreted by myeloid-derived suppressor cells (35). Potentially, the lung inflammatory environment inhibits effective containment of *K. pneumoniae*. However, the data are consistent with defective killing of initial founders and loss of a barrier for dissemination in *Nox2*^*-/-*^ mice, leading to higher lung founding populations, lung burdens, and populations in secondary organs that develop independently of growth in the lung. Nox2-dependent stresses may impose heterogeneity on bacterial populations in the lung, leading to uneven replication and metastatic dissemination.

The dynamics of bacteremia likely depend on the initial site of infection. Here, we used pneumonia as the primary infection due to its relevance in *K. pneumoniae* disease and death by AMR (9). The microenvironment of the lung during *K. pneumoniae* infection can change bacterial metabolism (36). Inflammatory reprogramming could change the bacterial ability to disseminate or influence fitness at secondary sites. For example, DsbA, involved in disulfide bond formation, is required for splenic fitness after tail vein injection but is dispensable when infection originates in the lung (32). Potentially during metastatic dissemination, bacteria that replicate in the lung are primed for increased splenic survival. In contrast, with direct dissemination bacteria that disseminate before replication may be less fit in the spleen. Further understanding of dissemination during Gram-negative bacteremia will integrate primary infections across other relevant sites.

Based on patient blood cultures, Gram-negative dissemination likely represents intermittent events of pathogen shedding rather than a single event or a continuous process. In the mouse model, intermittent shedding is supported by the high variability in measurements taken from the blood and is perhaps also reflected in the dissemination patterns observed here. Our study collected bacterial barcodes at a single timepoint, but additional resolution may be gained with measurements across time. Since higher lung CFU is linked to metastatic dissemination, this may be the dominant pattern observed at later timepoints once bacteria have undergone multiple rounds of division. Similarly, direct dissemination may be more likely at earlier time points. Importantly, both mechanisms were observed by 24 hours within each individual trial in our study. Further, in some cases individual mice displayed mixed patterns of dissemination to secondary organs. In *K. pneumoniae* lacking TatC and mice lacking Ccr2, animals displayed mixed dissemination modes to the liver and spleen, emphasizing some element of secondary organ tissue specificity in these findings. Future studies will investigate how the factors of infection route and timing influence dissemination patterns to secondary sites.

In summary, bacterial barcoding unveiled novel infection dynamics that were hidden when measuring only bacterial tissue burden. We discovered that patterns of heterogenous clonal expansion at primary sites underlie variability in dissemination and ultimately infection outcome. Quantifying the extent to which dissemination is metastatic or direct will substantially deepen our insights into how pathogens traverse throughout the host and contextualize how bacterial and host factors control infection outcomes. Our work demonstrates how high-complexity libraries of barcoded bacteria can be leveraged to reveal new paradigms of infection and better understand host-pathogen interactions during bacteremia.

## Supplemental Figures

**Supplemental Figure 1.**
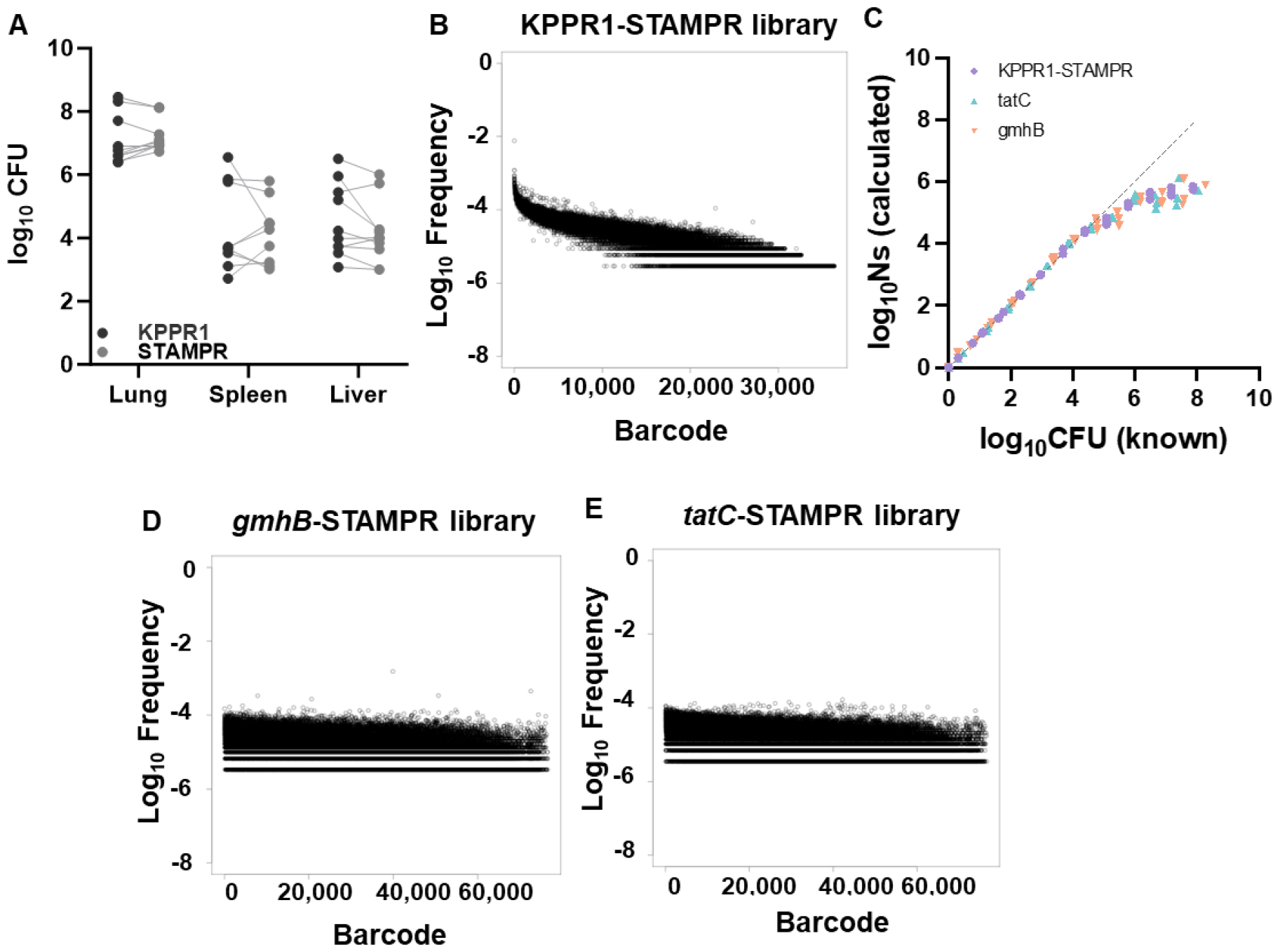
*K. pneumoniae* STAMPR libraries are fit and have even representation of barcodes. (A) Wild-type mice were co-infected at a 1:1 ratio with the unmarked KPPR1 strain and a single KPPR1-STAMPR barcoded clone in the murine model of pneumonia. At 24 hours, bacterial CFU was enumerated in each tissue and displayed as log_10_ CFU. Using paired *t*-tests, no significant difference in CFU abundance was detected between the KPPR1 or KPPR1-STAMPR strains. Barcoded *K. pneumoniae* libraries have a relatively even distribution of represented barcodes in the (B) KPPR1-STAMPR library. (C) A calibration curve with *in vitro* bottlenecks indicated that the *K. pneumoniae* barcoded libraries have a resolution limit of ~1x10^5^ clones. Barcoded *K. pneumoniae* libraries have a relatively even distribution of represented barcodes in the (D) *gmhB-*STAMPR, and (E) *tatC*-STAMPR libraries.

**Supplemental Figure 2.**
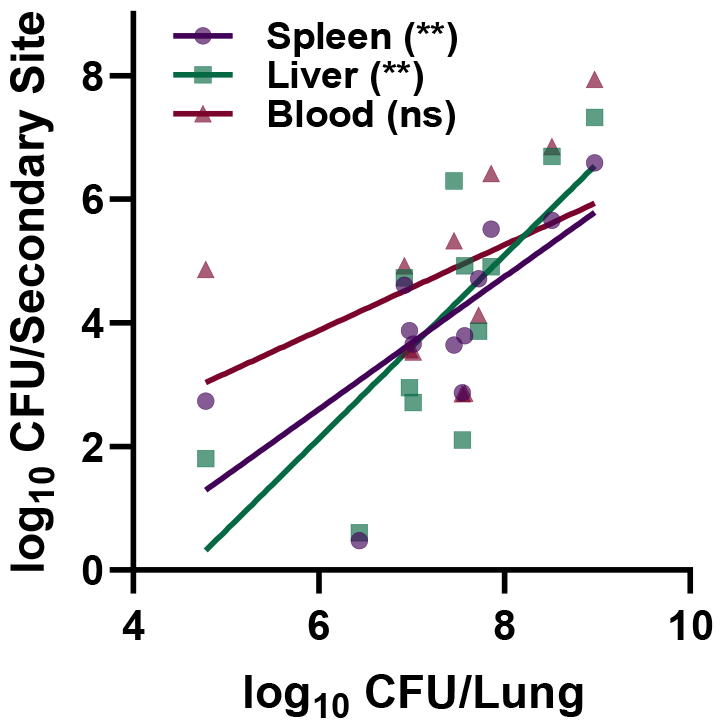
Lung bacterial burden correlated with CFU burden at secondary sites. The CFU data from Figure 1A was plotted to correlate CFU burden in the lung with burden in the spleen (purple circles), liver (green squares), and blood (pink triangles). Correlations were assessed using a Pearsons coefficient comparing lung CFU to the CFU at individual secondary sites, *\*\*p*<0.01.

**Supplemental Figure 3.**
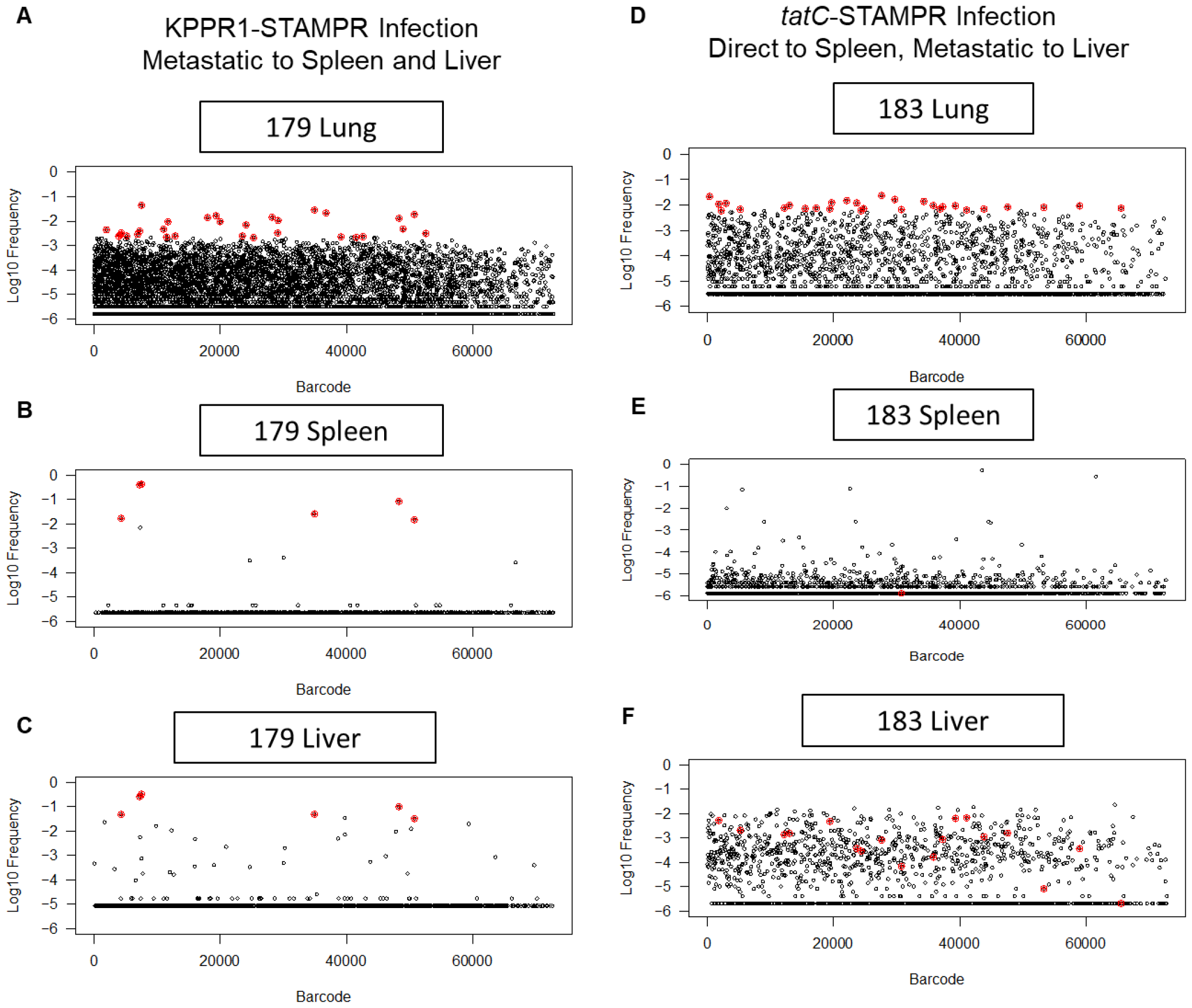
Representative frequency plots from a mouse demonstrating uniform dissemination patterns and one mouse demonstrating differential dissemination patterns to secondary organs. The STAMPR pipeline calculated the frequency of individual barcodes within the lung and spleen of two individual mice, one infected with KPPR1-STAMPR (A-C) and the other infected with *tatC-*STAMPR (D-F). Unique 25-nt barcodes were assigned random tags and plotted on the x-axis, the log_10_ frequency of each barcode within the indicated tissue is plotted on the y-axis. The most abundant barcodes within the lung for each mouse are highlighted in red, and indicated in the spleen if the barcode was also found at that site. In the wild-type bacterial infection, lung dissemination patterns were similar between the lung (A) and spleen (B) or the lung and liver (C), as demonstrated by a mouse displaying metastatic dissemination to each secondary site. A subset of mice infected with the *tatC-*STAMPR library demonstrated mixed dissemination patterns, as demonstrated by a mouse displaying direct dissemination from the lung (D) to the spleen (E) and metastatic dissemination from the lung to the liver (F).

## Methods and Materials

### Bacterial Strains and Reagents

All reagents were sourced from Sigma Aldrich unless noted otherwise. *K. pneumoniae* cultures were grown shaking overnight at 37°C in LB broth (Fisher Bioreagent, Ottowa, ON), and bacteria grown on LB plates were incubated at 30°C. Growth media was supplemented with 40μg/mL kanamycin (Sigma Aldrich, St. Louis, MO) to select for bacteria containing barcodes and/or with 50μg/mL hygromycin (Sigma-Aldrich) to select for knockout-strains generated by Lambda red mutagenesis. The bacterial strains used in this study are described in Supplemental Data 2, and primers used in the study are in Supplemental Data 3.

Lambda Red mutagenesis was used as previously described to generate *gmhB* (VK055_2352) and *tatC* (VK055_3142) isogenic knockouts containing a hygromycin resistance cassette (14, 30, 37). To generate electrocompetent KPPR1 harboring the pKD46 plasmid, an overnight culture was first grown at 30°C. The culture was then diluted into LB broth containing 50μg/mL spectinomycin, 50mM L-arabinose, 0.5mM EDTA (Promega, Madison, WI), and 10μM salicyclic acid and cultured until reaching exponential phase. The culture was placed on ice for 30 minutes, pelleted at 8,000xg for 15 minutes at 4°C, and serially washed with 50mL 1mM HEPES (pH 7.4; Gibco, Grand Island, NY), 50mL diH_2_O, and 20mL 10% glycerol. A hygromycin resistance cassette from the pSIM18 plasmid was amplified with primers containing 65 base pair homology at the 5’ end of the chromosomal site flanking the open reading frame of *gmhB* (14) or *tatC* (Supplemental Data 3). The purified fragment was electroporated into competent KPPR1-pKD46. Transformants were allowed to recover overnight, shaking at 30°C and then selected on agar with hygromycin. Each knockout was confirmed using primers flanking either *gmhB* or *tatC*.

### Construction of STAMPR Libraries

The MFDλpir-pSM1 plasmid donor library, containing random 30 nucleotide barcodes, was cultured in LB broth with kanamycin and 600μM DAP (15). To construct a *K. pneumoniae* library, a 1:1:1 ratio of the MFDλpir-pSM1 plasmid donor library, a helper donor (MFDλpir pLMP1039), and either wild-type KPPR1 (for KPPR1-STAMPR), Δ*gmhB*_hygro_ (for *gmhB-*STAMPR), or Δ*tatC*_hygro_ (for *tatC-*STAMPR) were spotted on a total of ten 0.45μm filters and incubated on plates containing 300μM DAP for 20-24 hours. After incubation, filters were combined and vigorously vortexed with PBS to release the transconjugants. The concentration of the recovered bacteria was determined by quantitative plating, and remaining transconjugants were plated onto multiple LB agar plates containing kanamycin and grown overnight at 30°C. All plates were pooled together by scraping recovered lawns into LB+25% glycerol and mixing thoroughly. The pools were split into multiple aliquots, labeled as the appropriate barcoded library, and stored at -80°C until further use.

### Murine Bacteremia

The study was approved by the University of Michigan Institutional Animal Care and Use Committee (protocol: PRO00011097), and performed with careful adherence to humane guidelines for animal handling (38). This study included male and female mice used between the ages of 8-12 weeks from the C57BL/6 lineage and included wild-type, *Nox2*^*-/-*^ (*Cybb*^*-/-*^), and *Ccr2*^*-/-*^ genotypes (32, 39, 40). All animals were bred and maintained at the University of Michigan or purchased directly (Jackson Laboratory, Bar Harbor, ME). Prior to each infection, *K. pneumoniae* overnight cultures for each strain were pelleted at 5,000xg for 15 minutes. The bacterial pellets were resuspended in PBS and the OD_600_ measured to adjust cultures to the desired concentration. For the pneumonia model, mice were anesthetized with isoflurane and 1x10^6^ *K. pneumoniae* CFU in 50μL PBS was administered retropharyngeally. Mice were sacrificed at 24 hours post-infection and lung, spleen, liver, and blood were collected. Blood samples were collected by cardiac punctures and dispensed into heparin coated tubes (BD, Franklin Lakes, NJ) to prevent clotting. Spleens were homogenized in 1mL PBS and livers were homogenized in 2mL PBS, and a 100μL sample was removed for quantitative plating to calculate the bacterial burden at each site. The remaining organ homogenate was plated onto LB dishes with kanamycin to recover the total number of barcodes within each site and plates were incubated overnight at 30°C. The following day, bacterial lawns were scraped into 10mL PBS+25% glycerol, keeping each mouse and tissue separate, and mixed thoroughly. 1x10^9^ CFU was removed from each sample and boiled to extract bacterial DNA. The remaining bacteria collected from lawn scraping were stored at -80°C.

### DNA Sequencing and STAMPR Analysis

The barcode-containing region was amplified from the bacterial DNA using the boiled bacterial cells in a PCR with OneTaq HS Quick-Load Master Mix (New England Biolabs) and custom primers (Supplemental Data 3). Primers contained TruSeq indexes and adapters for Illumina sequencing. PCR products were pooled, purified with a column kit (GeneJet PCR Purification Kit), and sequenced on either an Illumina MiSeq or Illumina NextSeq 1000.

Illumina sequencing read were demultiplexed with a custom R script, then trimmed and mapped to the appropriate reference library using either CLC Genomics Workbench (Qiagen) or custom R scripts. Mapped reads were exported as a CSV table. Any sample deemed to have low quality sequencing results, defined as <10,000 reads, was removed from the study and not included in further STAMPR pipeline analysis. An additional noise correction was performed as previously described (15) to adjust for increased index hopping in samples sequenced on the Illumina NextSeq. Founding population estimates were determined using STAMPR scripts (28) and genetic distance was estimated with Cavalli-Sforza cord distance as previously described (27).

N_s_ values were calculated as follows. After removing noise due to index hopping according to the STAMPR algorithm (28), the number of unique nonzero barcodes was determined. Then, for each output sample, the reference library was resampled according to a multinomial distribution with a sampling size equal to the total read count of the output sample. This resampled input vector was again resampled according to a multinomial distribution with varying sampling sizes. At each sampling size, the number of unique barcodes was calculated and a “reference resample curve” was created by plotting the sampling depth as the X-axis and the number of unique barcodes as the Y-axis. The number of unique barcodes in the output sample was used for inverse linear interpolation from the reference resample curve to calculate N_s_.

GD was calculated as the Cavalli-Sforza chord distance (27). FRD was calculated as follows. For two samples A and B, GD was first calculated (iteration 0). The most abundant barcode (geometric mean) in Samples A and B was removed, and GD was calculated again (iteration 1). Then, the first and second most abundant barcode were removed, and GD was calculated again (iteration 2). This procedure was repeated for 1000 iterations or until one sample had all barcodes >0 reads removed. The number of iterations that yield GD < 0.8, the threshold previously established for genetic relatedness, is defined as RD. Note that 0.8 is a relatively strict threshold, as samples with no overlapping barcodes have GD = (2√2)/π = 0.9. For pairs of samples where all 1000 iterations yielded GD < 0.8, RD was set to equal the number of unique barcodes with >0 reads in each sample. FRD(A-B) is defined as ln(RD(A-B) + 1) / ln(number of unique nonzero barcodes in sample B + 1).

Barcode counts and original scripts have been deposited on Github (https://github.com/hullahalli/stampr_rtisan). Individual heat maps for GD and FRD for each mouse in the study can be found in Supplemental Data 4.

### Statistical Analysis

At least two independent infections were performed for each group. Statistical significance was defined as a *p-*value<0.05 (GraphPad Prism) as determined using: ordinary one-way ANOVA with Dunnett’s multiple comparison to assess differences among multiple groups with one reference mean, ordinary one-way ANOVA with Tukey’s multiple comparison to assess differences among any groups, or unpaired *t-*tests to assess differences between two groups.

## References

1. Hajj J, Blaine N, Salavaci J, Jacoby D. The ‘Centrality of Sepsis‘: A Review on Incidence, Mortality, and Cost of Care. Healthcare (Basel). 2018;6(3).

2. Singer M, Deutschman CS, Seymour CW, Shankar-Hari M, Annane D, Bauer M, et al. The Third International Consensus Definitions for Sepsis and Septic Shock (Sepsis-3). JAMA. 2016;315(8):801–10.

3. Verway M, Brown KA, Marchand-Austin A, Diong C, Lee S, Langford B, et al. Prevalence and Mortality Associated with Bloodstream Organisms: a Population-Wide Retrospective Cohort Study. J Clin Microbiol. 2022;60(4):e0242921.

4. Wisplinghoff H, Bischoff T, Tallent SM, Seifert H, Wenzel RP, Edmond MB. Nosocomial bloodstream infections in US hospitals: analysis of 24,179 cases from a prospective nationwide surveillance study. Clin Infect Dis. 2004;39(3):309–17.

5. Marra AR, Camargo LF, Pignatari AC, Sukiennik T, Behar PR, Medeiros EA, et al. Nosocomial bloodstream infections in Brazilian hospitals: analysis of 2,563 cases from a prospective nationwide surveillance study. J Clin Microbiol. 2011;49(5):1866–71.

6. Diekema DJ, Hsueh PR, Mendes RE, Pfaller MA, Rolston KV, Sader HS, et al. The Microbiology of Bloodstream Infection: 20-Year Trends from the SENTRY Antimicrobial Surveillance Program. Antimicrob Agents Chemother. 2019;63(7):e00355–19.

7. CDC. Antibiotic Resistance Threats in the United State, 2013. Atlanta, GA: U.S. Department of Health and Human Services, Centers for Disease Control and Prevention; 2013.

8. CDC. Antibiotic Resistance Threats in the United States, 2019. Atlanta, GA: U.S. Department of Health and Human Services, Centers for Disease Control and Prevention; 2019.

9. Collaborators AR. Global burden of bacterial antimicrobial resistance in 2019: a systematic analysis. Lancet. 2022;399(10325):629–55.

10. Holmes CL, Anderson MT, Mobley HLT, Bachman MA. Pathogenesis of Gram-Negative Bacteremia. Clin Microbiol Rev. 2021;34(2).

11. Broadley SP, Plaumann A, Coletti R, Lehmann C, Wanisch A, Seidlmeier A, et al. Dual-Track Clearance of Circulating Bacteria Balances Rapid Restoration of Blood Sterility with Induction of Adaptive Immunity. Cell Host Microbe. 2016;20(1):36–48.

12. Milivojevic M, Dangeard AS, Kasper CA, Tschon T, Emmenlauer M, Pique C, et al. ALPK1 controls TIFA/TRAF6-dependent innate immunity against heptose-1,7-bisphosphate of gram-negative bacteria. PLoS Pathog. 2017;13(2):e1006224.

13. Zhou P, She Y, Dong N, Li P, He H, Borio A, et al. Alpha-kinase 1 is a cytosolic innate immune receptor for bacterial ADP-heptose. Nature. 2018;561(7721):122–6.

14. Holmes CL, Smith SN, Gurczynski SJ, Severin GB, Unverdorben LV, Vornhagen J, et al. The ADP-Heptose Biosynthesis Enzyme GmhB is a Conserved Gram-Negative Bacteremia Fitness Factor. Infect Immun. 2022;90(7):e0022422.

15. Fakoya B, Hullahalli K, Rubin DHF, Leitner DR, Chilengi R, Sack DA, et al. Nontoxigenic Vibrio cholerae Challenge Strains for Evaluating Vaccine Efficacy and Inferring Mechanisms of Protection. mBio. 2022;13(2):e0053922.

16. Hullahalli K, Dailey KG, Waldor MK. Innate immune responses yield tissue-specific bottlenecks that scale with pathogen dose. Proc Natl Acad Sci U S A. 2023;120(37):e2309151120.

17. Liu X, Kimmey JM, Matarazzo L, de Bakker V, Van Maele L, Sirard JC, et al. Exploration of Bacterial Bottlenecks and Streptococcus pneumoniae Pathogenesis by CRISPRi-Seq. Cell Host Microbe. 2021;29(1):107-20.e6.

18. Wincott CJ, Sritharan G, Benns HJ, May D, Gilabert-Carbajo C, Bunyan M, et al. Cellular barcoding of protozoan pathogens reveals the within-host population dynamics of Toxoplasma gondii host colonization. Cell Rep Methods. 2022;2(8):100274.

19. Aggarwal SD, Lees JA, Jacobs NT, Bee GCW, Abruzzo AR, Weiser JN. BlpC-mediated selfish program leads to rapid loss of Streptococcus pneumoniae clonal diversity during infection. Cell Host Microbe. 2023;31(1):124-34.e5.

20. Woodward SE, Vogt SL, Peña-Díaz J, Melnyk RA, Cirstea M, Serapio-Palacios A, et al. Gastric acid and escape to systemic circulation represent major bottlenecks to host infection by Citrobacter rodentium. ISME J. 2023;17(1):36–46.

21. Campbell IW, Hullahalli K, Turner JR, Waldor MK. Quantitative dose-response analysis untangles host bottlenecks to enteric infection. Nat Commun. 2023;14(1):456.

22. Zhang T, Sasabe J, Hullahalli K, Sit B, Waldor MK. Increased Listeria monocytogenes Dissemination and Altered Population Dynamics in Muc2-Deficient Mice. Infect Immun. 2021;89(4).

23. Hullahalli K, Waldor MK. Pathogen clonal expansion underlies multiorgan dissemination and organ-specific outcomes during murine systemic infection. Elife. 2021;10.

24. Mahmutovic A, Gillman AN, Lauksund S, Robson Moe NA, Manzi A, Storflor M, et al. RESTAMP-Rate estimates by sequence-tag analysis of microbial populations. Comput Struct Biotechnol J. 2021;19:1035–51.

25. Bachta KER, Allen JP, Cheung BH, Chiu CH, Hauser AR. Systemic infection facilitates transmission of Pseudomonas aeruginosa in mice. Nat Commun. 2020;11(1):543.

26. Zhang T, Abel S, Abel Zur Wiesch P, Sasabe J, Davis BM, Higgins DE, et al. Deciphering the landscape of host barriers to Listeria monocytogenes infection. Proc Natl Acad Sci U S A. 2017;114(24):6334–9.

27. Abel S, Abel zur Wiesch P, Chang HH, Davis BM, Lipsitch M, Waldor MK. Sequence tag-based analysis of microbial population dynamics. Nat Methods. 2015;12(3):223–6, 3 p following 6.

28. Hullahalli K, Pritchard JR, Waldor MK. Refined Quantification of Infection Bottlenecks and Pathogen Dissemination with STAMPR. mSystems. 2021;6(4):e0088721.

29. Broberg CA, Wu W, Cavalcoli JD, Miller VL, Bachman MA. Complete Genome Sequence of Klebsiella pneumoniae Strain ATCC 43816 KPPR1, a Rifampin-Resistant Mutant Commonly Used in Animal, Genetic, and Molecular Biology Studies. Genome Announc. 2014;2(5).

30. Bachman MA, Breen P, Deornellas V, Mu Q, Zhao L, Wu W, et al. Genome-Wide Identification of Klebsiella pneumoniae Fitness Genes during Lung Infection. mBio. 2015;6(3):e00775.

31. Anderson MT, Mitchell LA, Zhao L, Mobley HLT. Citrobacter freundii fitness during bloodstream infection. Sci Rep. 2018;8(1):11792.

32. Holmes CL, Wilcox AE, Forsyth V, Smith SN, Moricz BS, Unverdorben LV, et al. Klebsiella pneumoniae causes bacteremia using factors that mediate tissue-specific fitness and resistance to oxidative stress. PLoS Pathog. 2023;19(7):e1011233.

33. Mobley HLT, Anderson MT, Moricz BS, Severin GB, Holmes CL, Ottosen EN, et al. Fitness Factor Genes Conserved within the Multi-species Core Genome of Gram-negative Enterobacterales Species Contribute to Bacteremia Pathogenesis. BioRxiv [Preprin], doi: 101101/20240318585282v1. 2024.

34. Garcia-Weber D, Arrieumerlou C. ADP-heptose: a bacterial PAMP detected by the host sensor ALPK1. Cell Mol Life Sci. 2021;78(1):17–29.

35. Ahn D, Peñaloza H, Wang Z, Wickersham M, Parker D, Patel P, et al. Acquired resistance to innate immune clearance promotes Klebsiella pneumoniae ST258 pulmonary infection. JCI Insight. 2016;1(17):e89704.

36. Wong Fok Lung T, Charytonowicz D, Beaumont KG, Shah SS, Sridhar SH, Gorrie CL, et al. Klebsiella pneumoniae induces host metabolic stress that promotes tolerance to pulmonary infection. Cell Metab. 2022;34(5):761-74.e9.

37. Datsenko KA, Wanner BL. One-step inactivation of chromosomal genes in Escherichia coli K-12 using PCR products. Proc Natl Acad Sci U S A. 2000;97(12):6640–5.

38. (US) NRC. Guide for the care and use of laboratory animals. Washington, D.C.: National Academies Press; 2011.

39. Moore BB, Paine R, Christensen PJ, Moore TA, Sitterding S, Ngan R, et al. Protection from pulmonary fibrosis in the absence of CCR2 signaling. J Immunol. 2001;167(8):4368–77.

40. Werner JL, Escolero SG, Hewlett JT, Mak TN, Williams BP, Eishi Y, et al. Induction of Pulmonary Granuloma Formation by Propionibacterium acnes Is Regulated by MyD88 and Nox2. Am J Respir Cell Mol Biol. 2017;56(1):121–30.

