## Supplemental Data 4 for "Patterns of *Klebsiella pneumoniae* bacteremic dissemination from the lung"

**GD** **FRD**

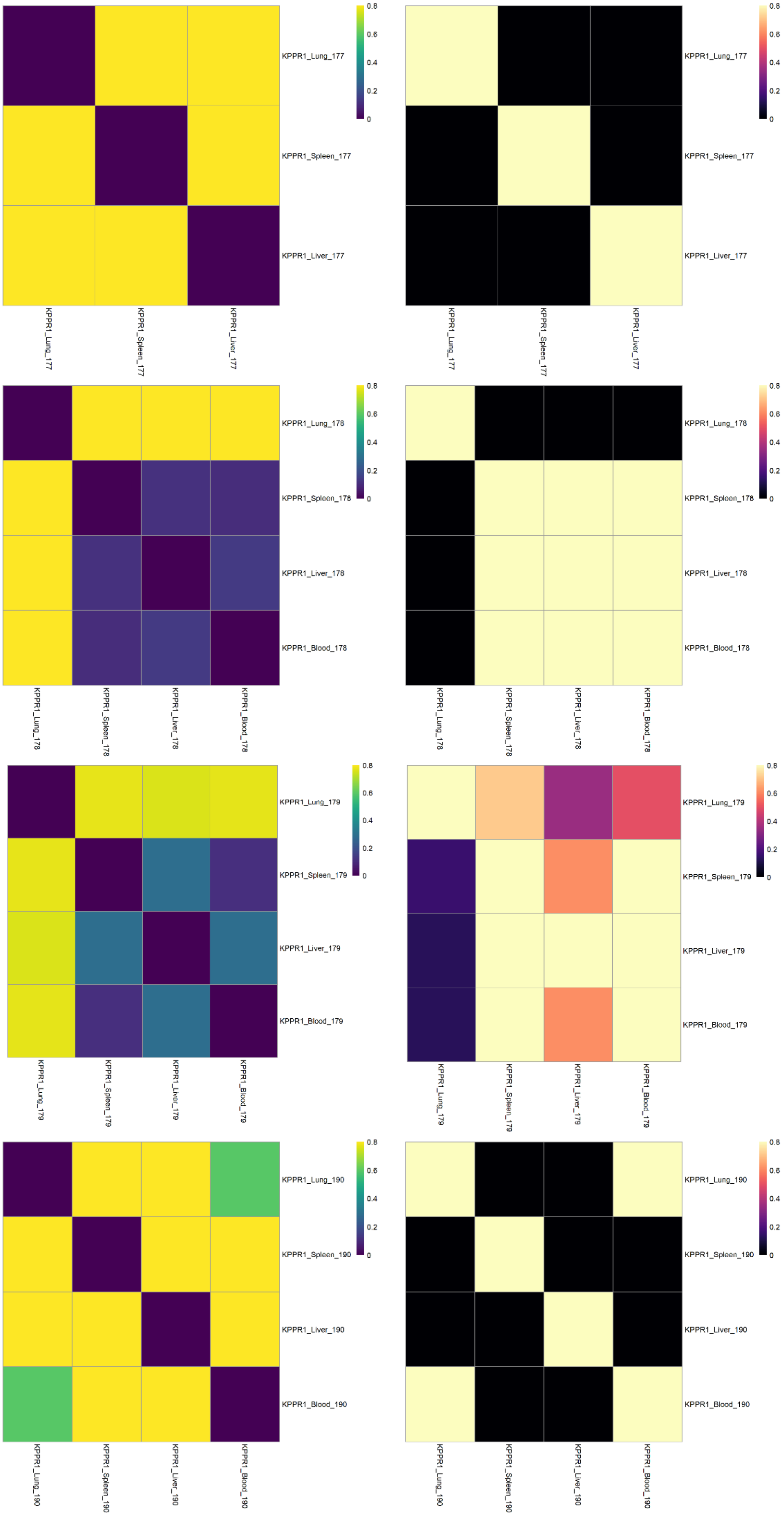


**Notes**

Mouse 177

no blood CFU recovered

KPPR1

Mouse 178

KPPR1

Mouse 179

KPPR1

Mouse 190

KPPR1

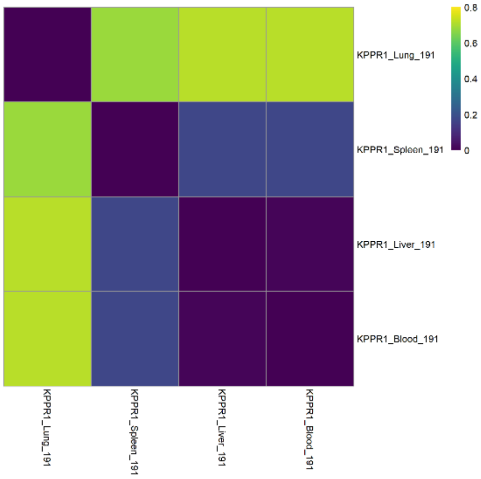


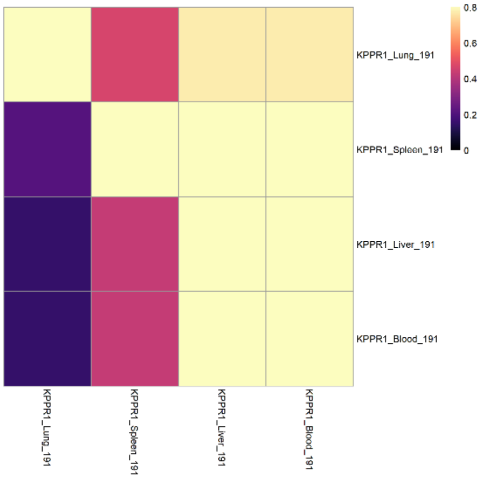


Mouse 191

KPPR1

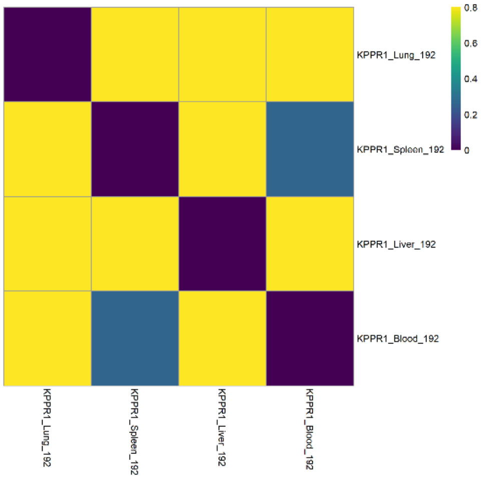


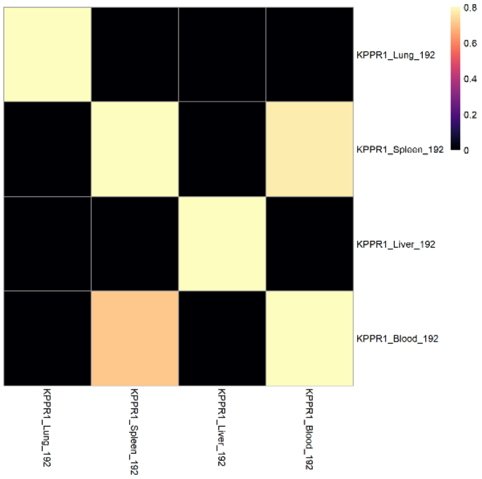


Mouse 192

KPPR1

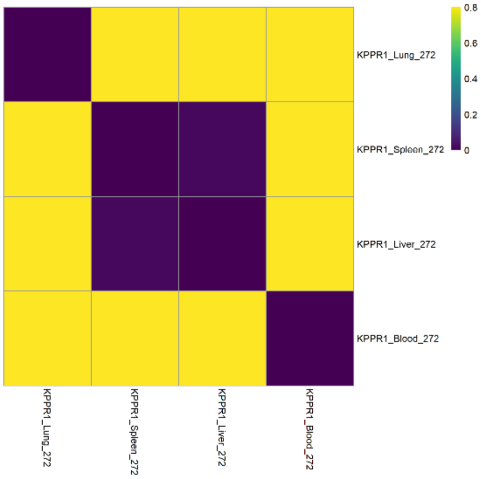


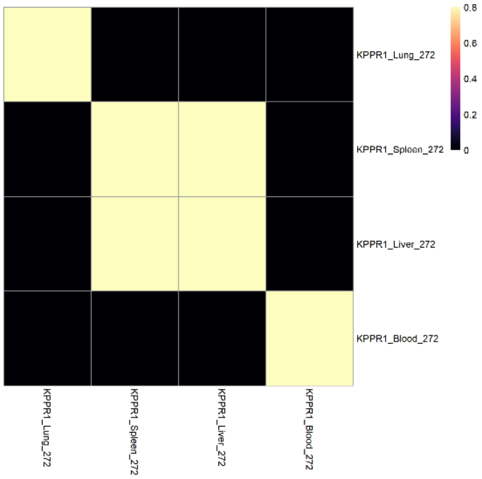


Mouse 272

Blood sample removed from downstream STAMPR analysis due to low seuqncing quality

KPPR1

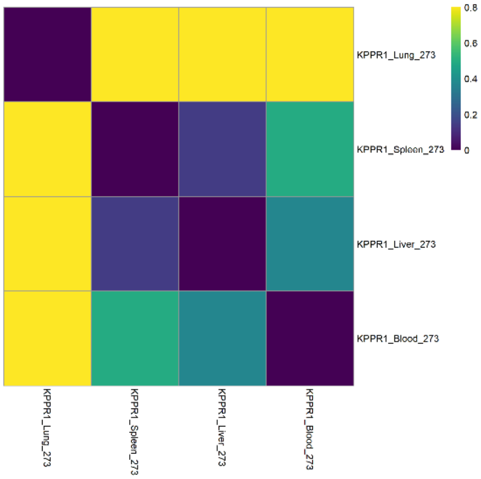


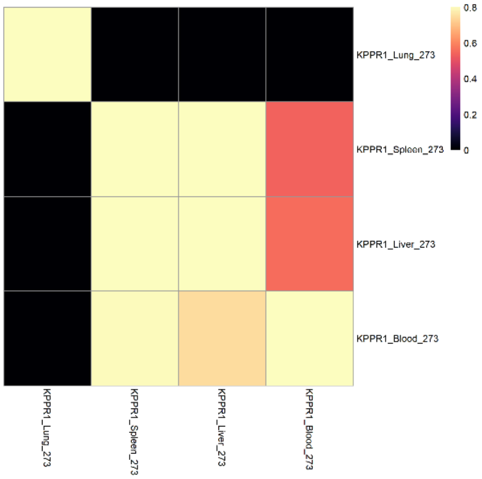


Mouse 273

KPPR1

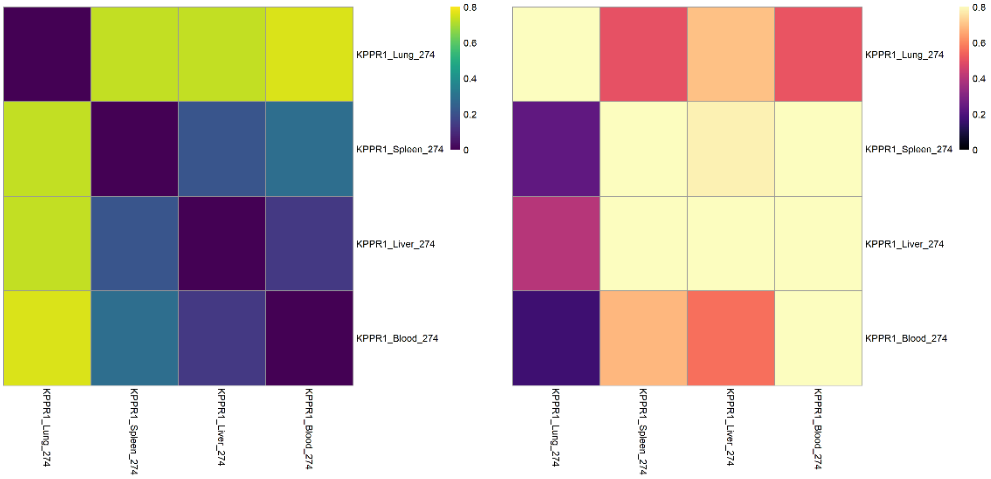


Mouse 274

KPPR1

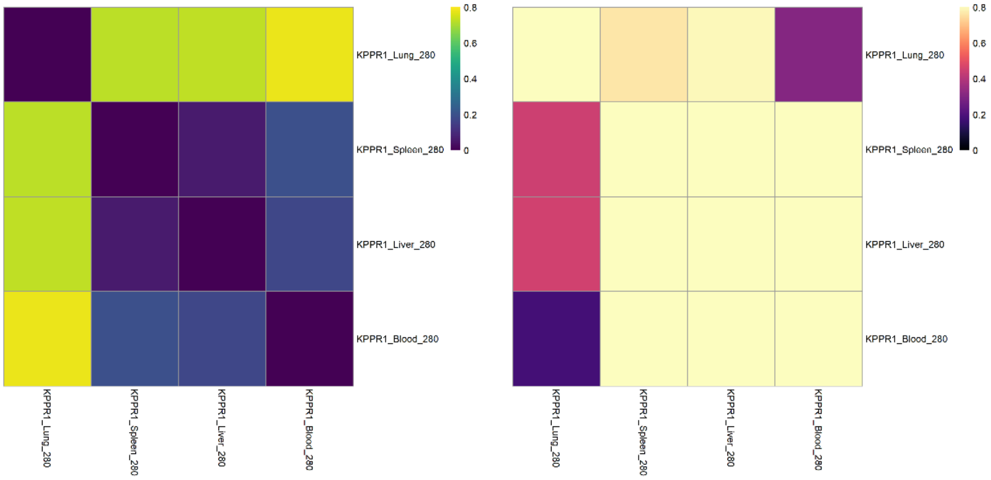


Mouse 280

KPPR1

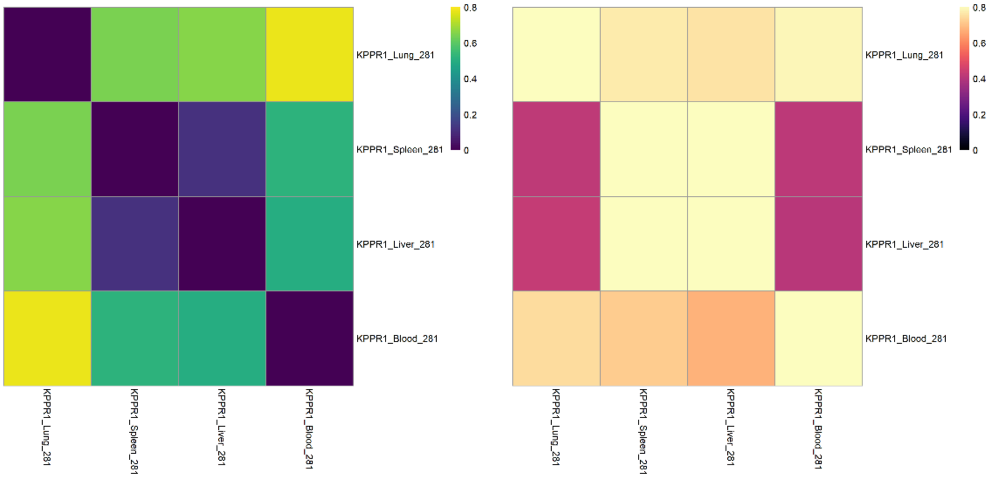


Mouse 281

KPPR1

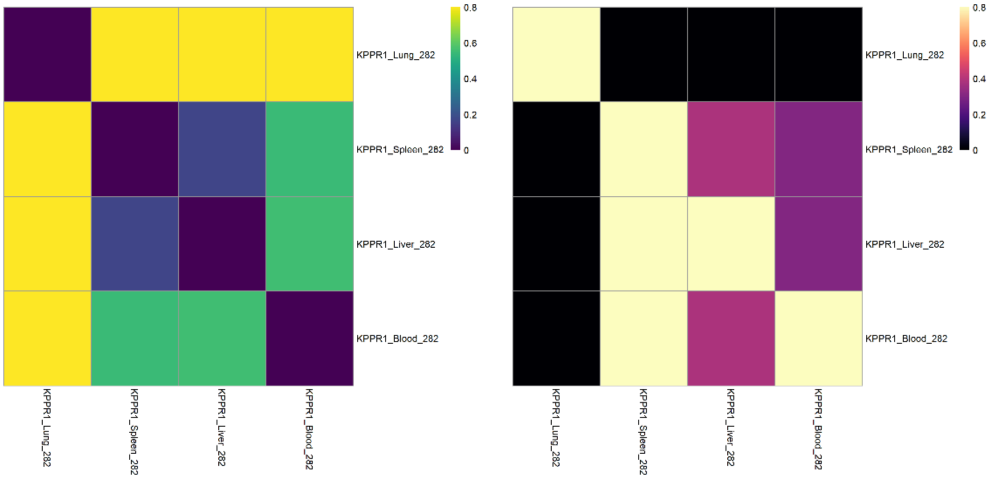


Mouse 282

Liver sample removed from downstream STAMPR analysis due to low seuqncing quality

KPPR1

Mouse 181

*gmhB*

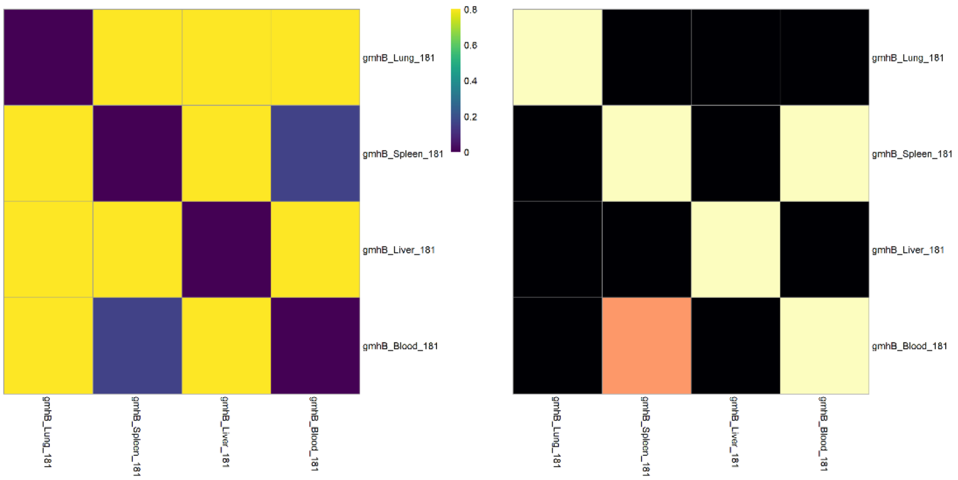


Mouse 182

*gmhB*

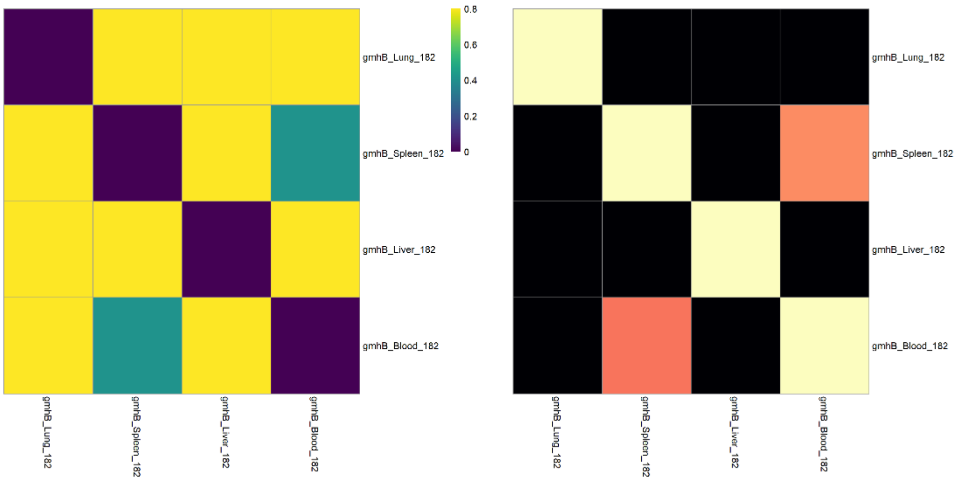


Mouse 194

*gmhB*

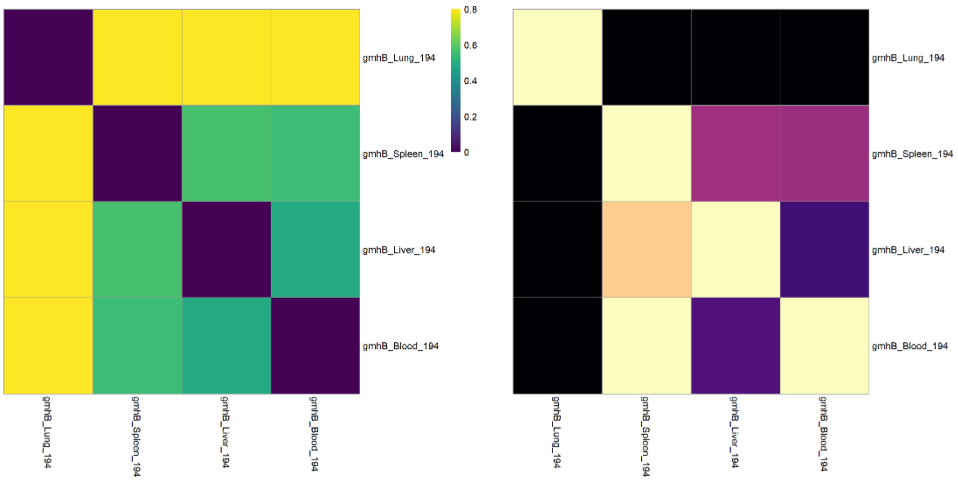


Mouse 195

*gmhB*

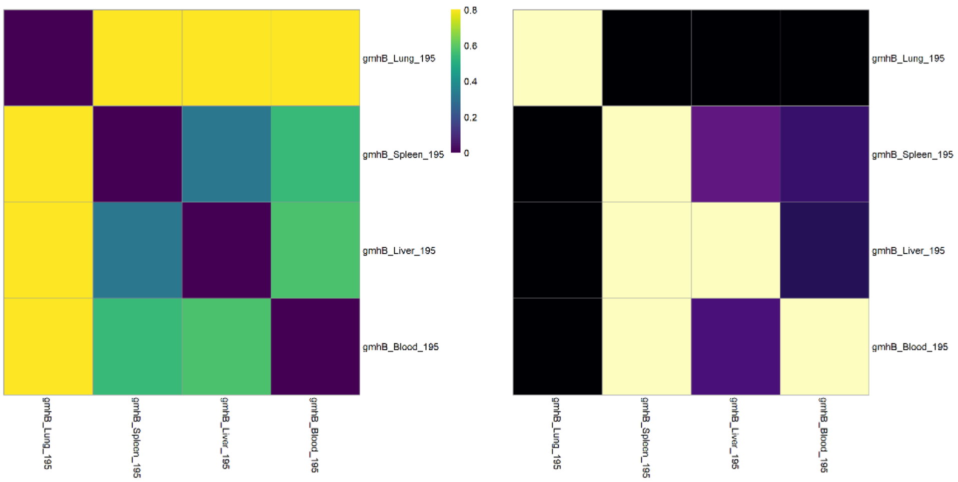

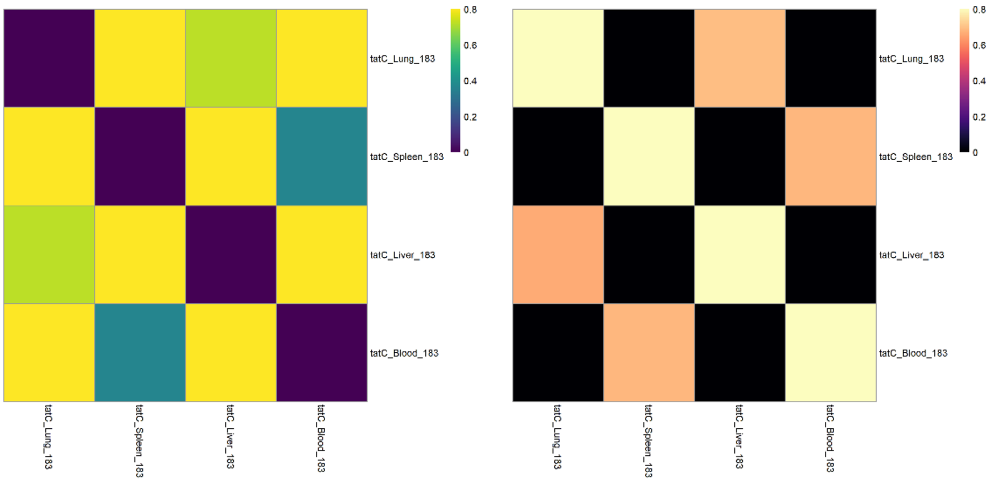


Mouse 183

*tatC*

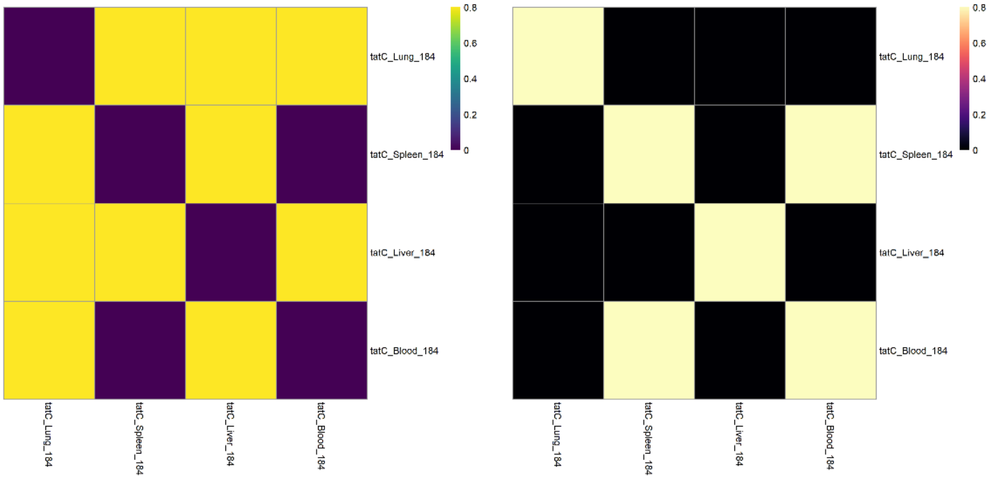


Mouse 184

*tatC*

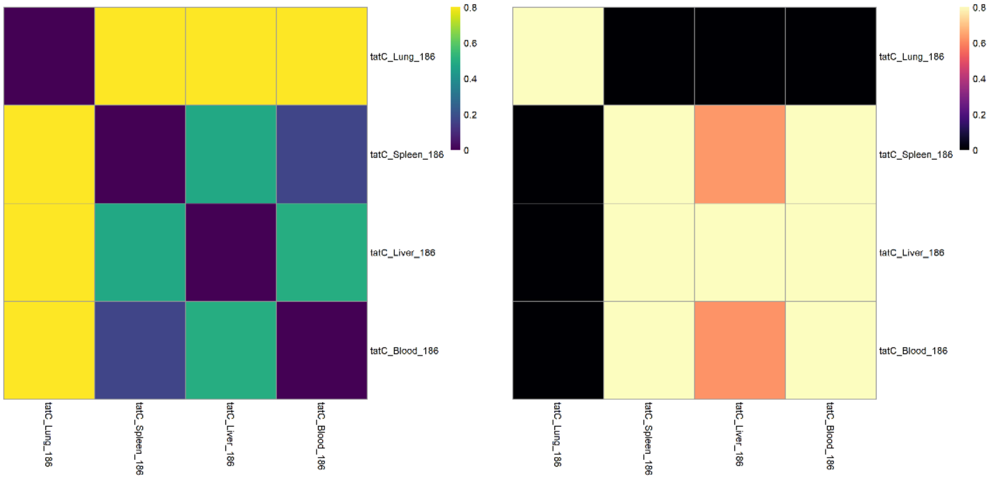


Mouse 186

*tatC*

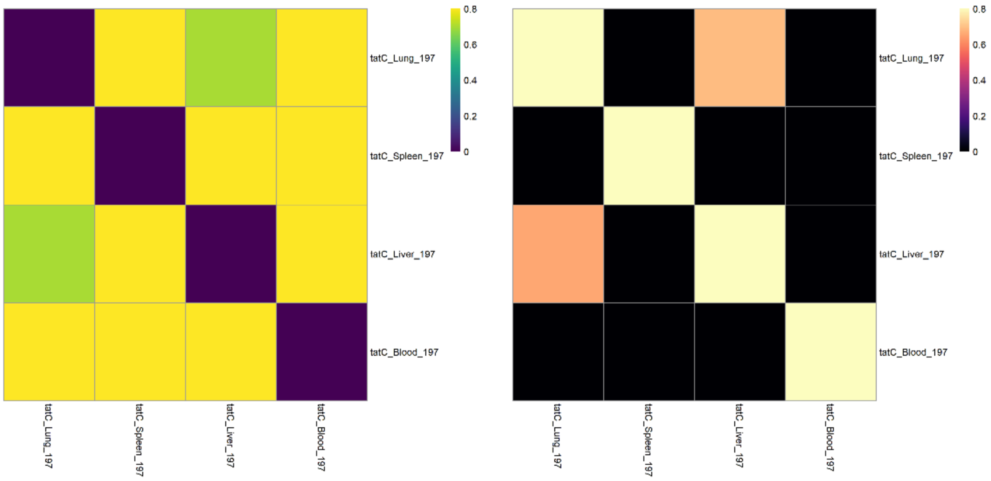


Mouse 197

*tatC*

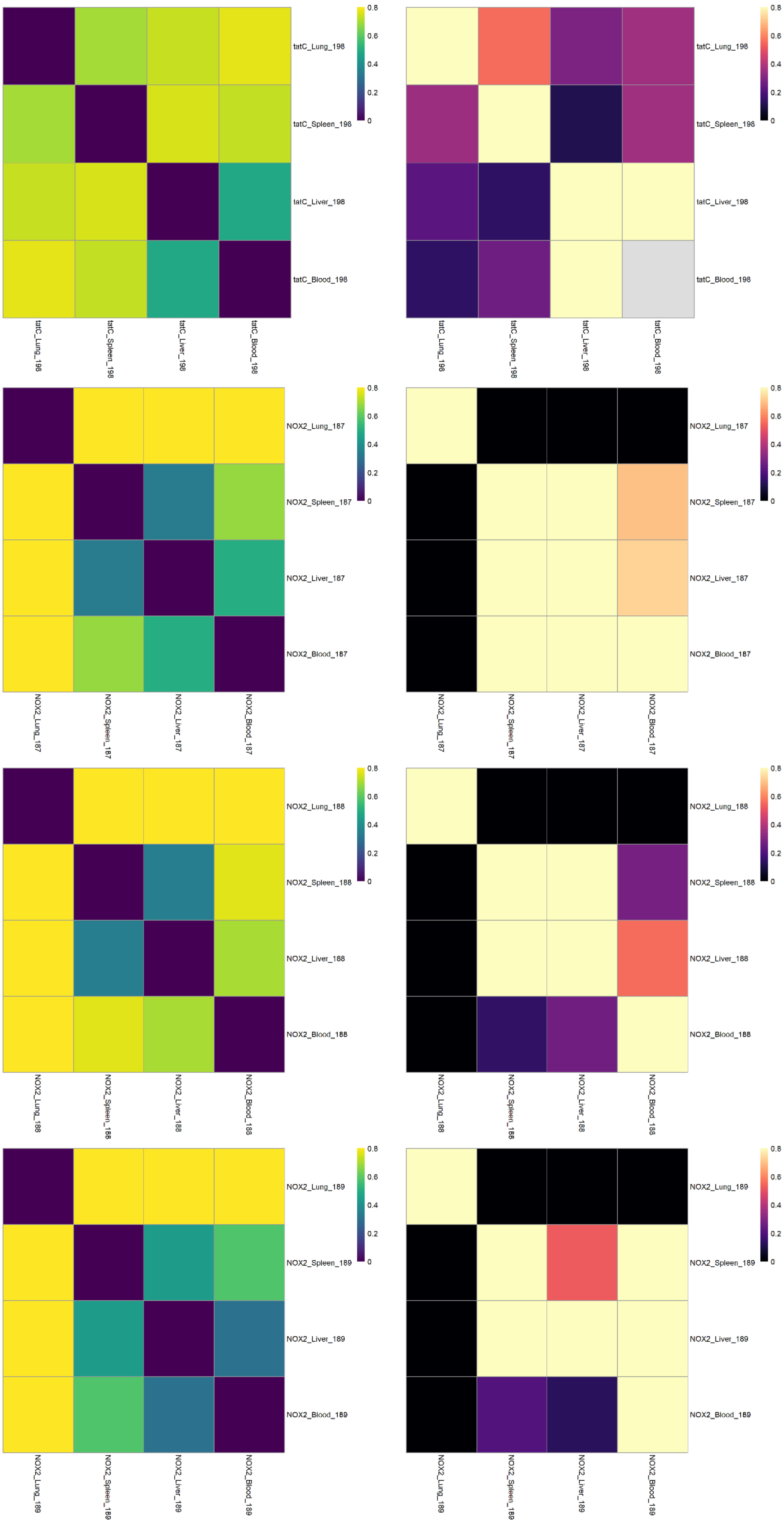


Mouse 198

*tatC*

Mouse 187

*Nox2*$^{-/-}$

Mouse 188

*Nox2*$^{-/-}$

Mouse 189

*Nox2*$^{-/-}$

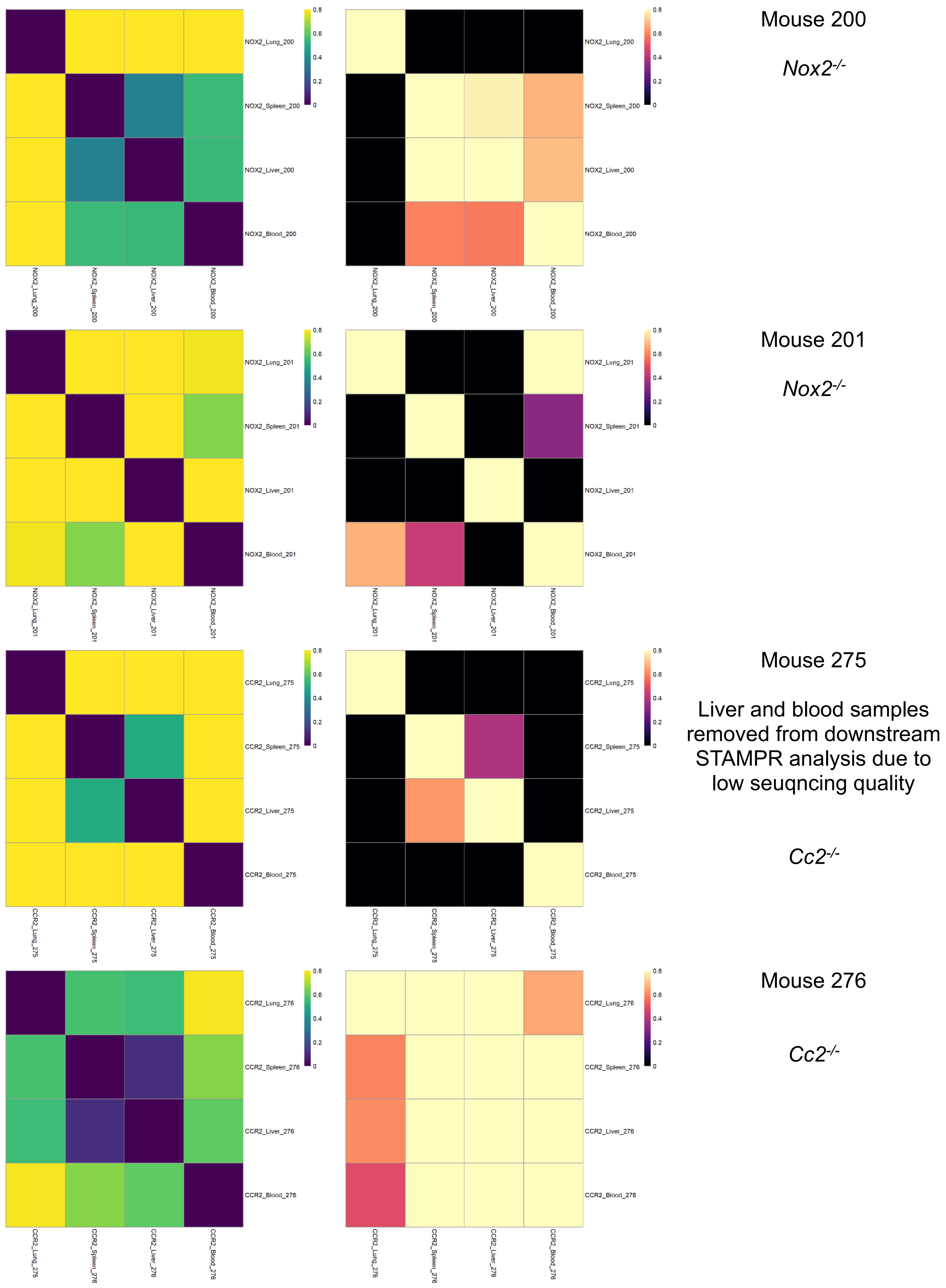

Mouse 200
Nox2-/-
NOX2_Lung_200
NOX2_Spleen_200
NOX2_Liver_200
NOX2_Blood_200
Mouse 201
Nox2-/-
NOX2_Lung_201
NOX2_Spleen_201
NOX2_Liver_201
NOX2_Blood_201
Mouse 275
Liver and blood samples removed from downstream STAMPR analysis due to low seuqncing quality
Cc2-/-
CCR2_Lung_275
CCR2_Spleen_275
CCR2_Liver_275
CCR2_Blood_275
Mouse 276
Cc2-/-
CCR2_Lung_276
CCR2_Spleen_276
CCR2_Liver_276
CCR2_Blood_276
0
0.2
0.4
0.6
0.8

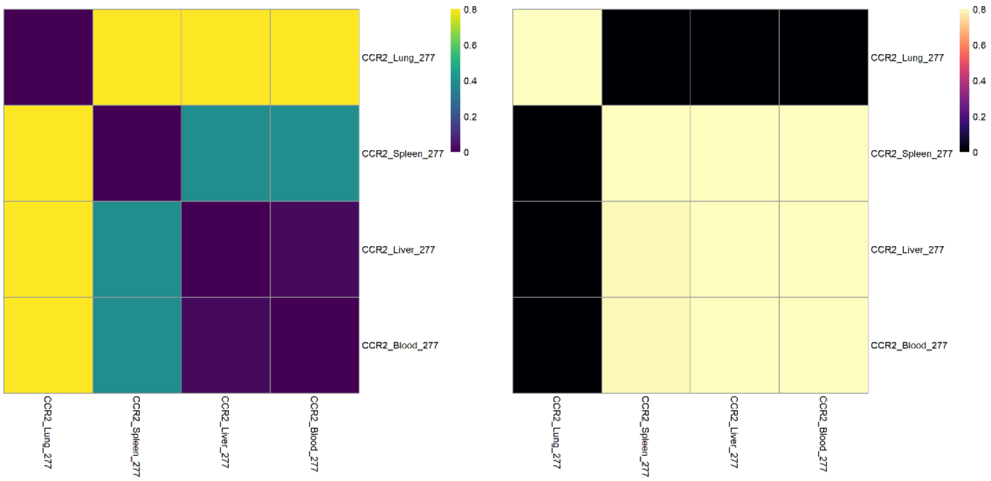


Mouse 277

*Cc2*$^{-/-}$

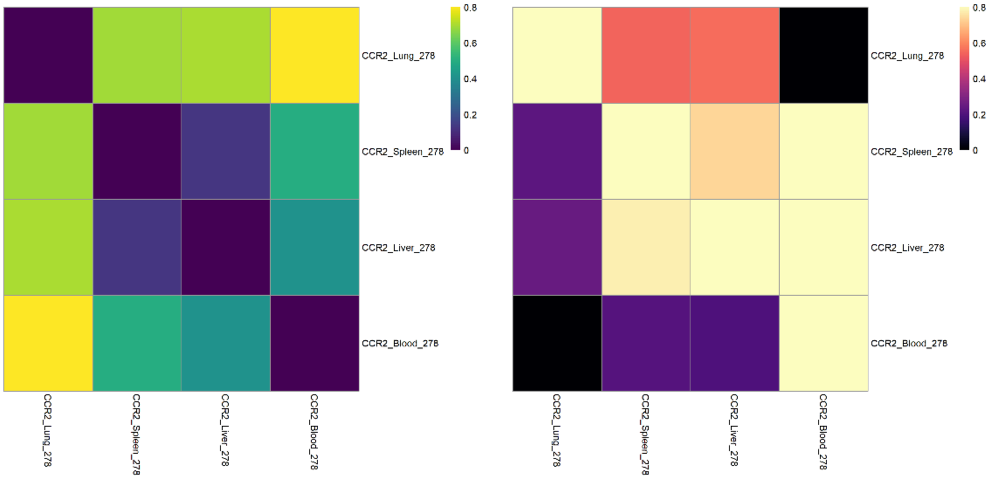


Mouse 278

*Cc2*$^{-/-}$

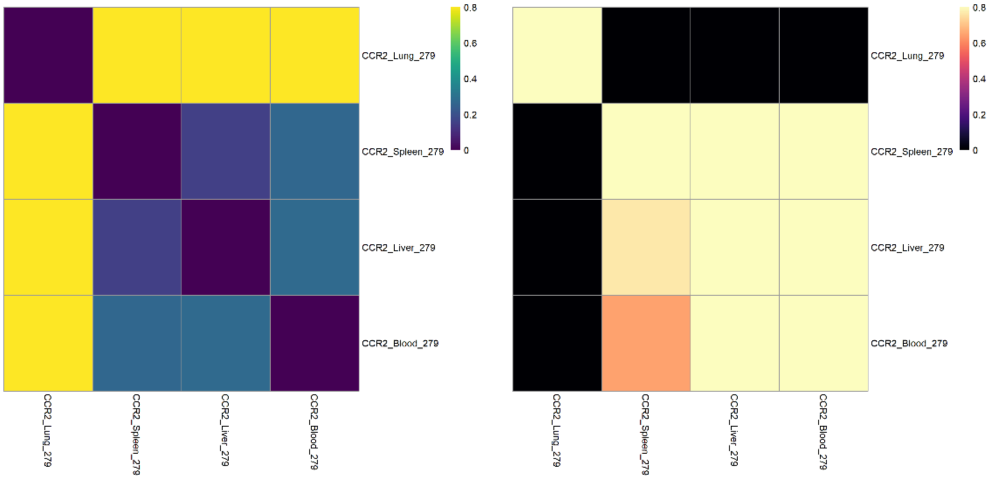


Mouse 279

*Cc2*$^{-/-}$

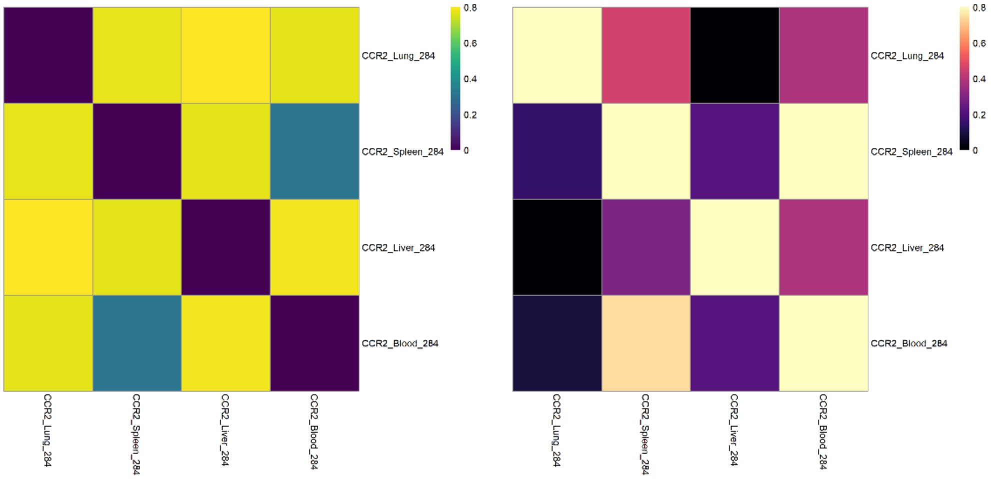


Mouse 284

Liver sample removed from downstream STAMPR analysis due to low seuqncing quality

*Cc2*$^{-/-}$

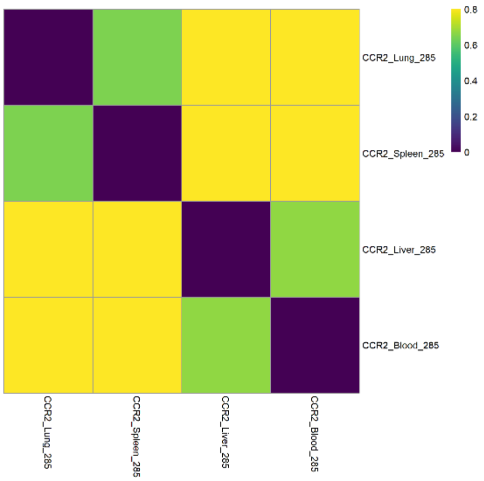


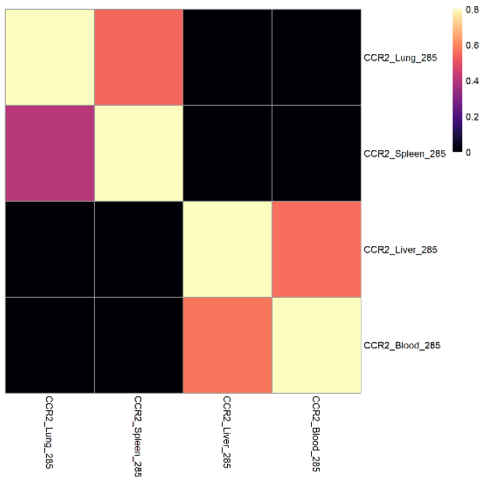


Mouse 285

Liver and blood samples removed from downstream STAMPR analysis due to low seuqncing quality

*Cc2*$^{-/-}$

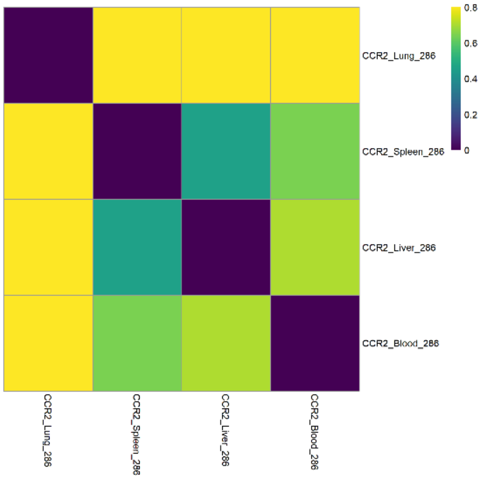


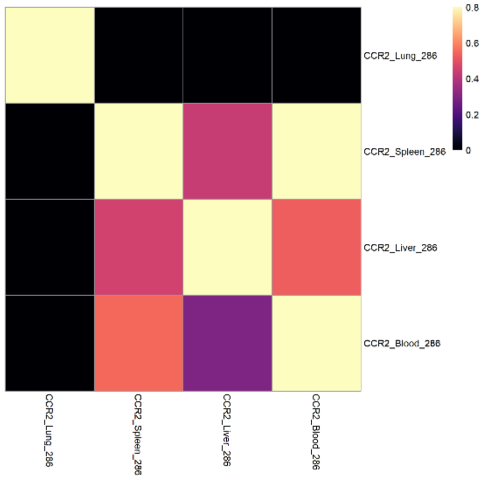


Mouse 286

Liver sample removed from downstream STAMPR analysis due to low seuqncing quality

*Cc2*$^{-/-}$
